# Biochemical characterization of recombinant UDP-sugar pyrophosphorylase and galactinol synthase from *Brachypodium distachyon*

**DOI:** 10.1101/2020.07.21.212928

**Authors:** Romina I. Minen, María P. Martínez, Alberto A. Iglesias, Carlos M. Figueroa

## Abstract

Raffinose (Raf) protects plant cells during seed desiccation and under different abiotic stress conditions. The biosynthesis of Raf starts with the production of UDP-galactose by UDP-sugar pyrophosphorylase (USPPase) and continues with the synthesis of galactinol by galactinol synthase (GolSase). Galactinol is then used by Raf synthase to produce Raf. In this work, we report the biochemical characterization of USPPase (*Bdi*USPPase) and GolSase 1 (*Bdi*GolSase1) from *Brachypodium distachyon*. The catalytic efficiency of *Bdi*USPPase was similar with galactose 1-phosphate and glucose 1-phosphate, but 5-to 17-fold lower with other sugar 1-phosphates. The catalytic efficiency of *Bdi*GolSase1 with UDP-galactose was three orders of magnitude higher than with UDP-glucose. A structural model of *Bdi*GolSase1 allowed us to determine the residues putatively involved in the binding of substrates. Among these, we found that Cys^261^ lies within the putative catalytic pocket. *Bdi*GolSase1 was inactivated by oxidation with diamide and H_2_O_2_. The activity of the diamide-oxidized enzyme was recovered by reduction with dithiothreitol or *E. coli* thioredoxin, suggesting that *Bdi*GolSase1 is redox-regulated.

## 1. Introduction

Raffinose (Raf) is an α-1,6-galactosyl extension of sucrose (Suc). Raf metabolism seems to be universal in angiosperms, but oligosaccharides derived from Raf (such as stachyose and verbascose) are only found in certain plant species (Handley *et al*., 1983; Peters and Keller, 2009). Raf is involved in membrane stabilization during seed desiccation (Downie *et al*., 2003) and used for carbon transport to heterotrophic tissues in plants from the Cucurbitaceae family (Haritatos *et al*., 1996). Raf also accumulates in leaves of plants exposed to several abiotic stress conditions, including heat, cold, salinity, and drought (Bachmann *et al*., 1994; Taji *et al*., 2002; Panikulangara, 2004; Zuther *et al*., 2004; Peters and Keller, 2009).

The first step in the pathway of Raf synthesis is the production of UDP-galactose (UDP-Gal), which is not specific to this route but provides an essential substrate. Plants have two alternative ways to produce this sugar nucleotide (Decker and Kleczkowski, 2019). One possibility is the Leloir pathway (Leloir, 1951), which involves the reversible conversion of UDP-glucose (UDP-Glc) to UDP-Gal by UDP-Glc 4-epimerase (EC 5.1.3.2). Alternatively, UDP-Gal can be synthesized from galactose 1-phosphate (Gal1P) and UTP, in a reaction catalyzed by UDP-sugar pyrophosphorylase (USPPase, EC 2.7.7.64) (Dai, 2006). Previous studies have shown that higher plants appear to lack, or have a limited activity of, the critical uridylyltransferase enzyme of the Leloir-pathway (Frey, 1996; Dai, 2006), suggesting that a USPPase is responsible for Gal1P metabolism (Smart and Pharr, 1981; Studer Feusi *et al*., 1999; Decker and Kleczkowski, 2019). In the model plant *Arabidopsis thaliana*, USPPase is expressed in various tissues and developmental stages (Kotake *et al*., 2007), and mutation of the gene coding for this enzyme leads to abnormal pollen production (Schnurr *et al*., 2006). USPPases characterized so far show relaxed specificity towards the phosphorylated sugar used as a substrate. For instance, USPPases from *A. thaliana* and barley show activity with Gal1P, glucose 1-phosphate (Glc1P), and glucuronic acid 1-phosphate (GlcA1P), among others (Decker and Kleczkowski, 2017; Wahl *et al*., 2017).

The route of Raf synthesis continues with galactinol synthase (GolSase, EC 2.4.1.123), which produces galactinol from UDP-Gal and *myo*-inositol. Raf synthase (EC 2.4.1.82) then transfers the galactosyl moiety from galactinol to a preformed Suc molecule, thus producing Raf (Lehle and Tanner, 1973; Castillo, 1990; Liu *et al*., 1995; Peterbauer *et al*., 2002; Zhao *et al*., 2004; Schneider and Keller, 2009; Wang *et al*., 2009; Sengupta *et al*., 2015). GolSase is a member of the glycosyltransferase family 8 (GT8), which includes many enzymes involved in the biosynthesis of various sugar conjugates (Sengupta *et al*., 2012; Dos Santos *et al*., 2015; Yin *et al*., 2018). GolSase catalyzes the first committed step in Raf synthesis; therefore, it plays a crucial role in carbon partitioning between Suc and Raf (Robbins and Pharr, 1987; Saravitz *et al*., 1987; Smith *et al*., 1991; Hitz *et al*., 2002).

The expression of genes coding for GolSase has been studied in several plant species exposed to various types of stressful conditions. In *A. thaliana*, expression of *AtGolS1* and *AtGolS2* is induced by drought, heat, and high salinity, while expression of *AtGolS3* is triggered by cold. Increased expression of *GolSase* genes has been described in other species (like *Populus trichocarpa, Coffea canephora, Cucumis melo, Solanum lycopersicum*, and *Oryza sativa*) exposed to drought and cold. Overexpression of the *AtGolS2* gene in both *A. thaliana* and *O. sativa* produced plants with higher levels of galactinol and Raf, which in turn increased drought tolerance (Taji *et al*., 2002; Volk *et al*., 2003; Saito and Yoshida, 2011; Zhou *et al*., 2014; Pluskota *et al*., 2015; Shimosaka and Ozawa, 2015; de Souza Vidigal *et al*., 2016; Selvaraj *et al*., 2017). Considering the importance of GolSase for Raf metabolism, little is known about this enzyme’s kinetic and structural properties. Most reports are characterized by containing scarce, incomplete kinetic determinations and all the work performed so far has been done with GolSases from dicot species (Handley and Pharr, 1982; Handley *et al*., 1983; Smith *et al*., 1991; Liu *et al*., 1995).

*Brachypodium distachyon* (Poaceae family) has many features that make it an excellent model organism for research. Concerning evolution, *B. distachyon* is related to the major cereal grain species, such as wheat, barley, oat, rice, and rye (Vogel and Hill, 2008; Vogel *et al*., 2010; Filiz *et al*., 2015). Additionally, several works deal with the effects triggered by different types of abiotic stress conditions (such as salinity, drought, cold and heat) on this plant species, although most of them are focused on the consequences at the transcript-level (Li *et al*., 2012; Verelst *et al*., 2013; Lv *et al*., 2014; Priest *et al*., 2014). It has been shown that *B. distachyon* produces galactinol and Raf when exposed to cold and drought (Ryu *et al*., 2014; Filiz *et al*., 2015; Ahkami *et al*., 2019). In this work, we recombinantly produced, purified, and characterized *B. distachyon* USPPase (*Bdi*USPPase) and GolSase 1 (*Bdi*GolSase1). Our results show that *Bdi*USPPase could provide UDP-Gal for galactinol synthesis by *Bdi*GolSase1. We also obtained a 3D structural model of *Bdi*GolSase1, which allowed us to identify residues putatively involved in catalysis and redox regulation. Finally, we explored the effect of different oxidizing and reducing agents on the activity of *Bdi*GolSase1.

## 2. Materials and methods

### 2.1. Bacterial strains and reagents

*Escherichia coli* TOP10 cells (Invitrogen) were used for cloning procedures and plasmid maintenance. Protein expression was carried out using *E. coli* BL21 Star (DE3) (Invitrogen). The substrates used to determine enzyme activity were from Sigma-Aldrich. All other reagents were of the highest purity available.

### 2.2. Phylogenetic analysis

The protein sequence of *B. distachyon* GolSase 1 was downloaded from the NCBI database (https://www.ncbi.nlm.nih.gov/). Using this sequence as a query, we obtained 95 plant sequences from Phytozome v12.1 (http://www.phytozome.org/). Sequences were analyzed with the program BioEdit 7.0.5.3 (Hall, 1999) and aligned in the ClustalW server (http://www.genome.jp/tools/clustalw/). Sequence codes are presented in Supplemental Table S1, and the alignment used for tree reconstruction is available in Supplemental File S1. The phylogenetic tree was reconstructed in Seaview 5.0.1 (Gouy *et al*., 2010) with the Neighbor-Joining method, with a bootstrap of 1,000. The tree was prepared in the FigTree 1.4.4 program (http://tree.bio.ed.ac.uk/).

### 2.3. *Synthesis of the genes encoding* Bdi*USPPase and* Bdi*GolSase1*

The sequences encoding *Bdi*USPPase (NCBI accession number KQK17397.1) and *Bdi*GolSase1 (NCBI accession number KQK21944.1) were *de novo* synthesized by Bio Basic Inc. (Canada) and inserted into the pUC57 vector. The genes were subcloned into the pET28c vector (Novagen) between the *Nde*I and *Sac*I sites to obtain the recombinant proteins fused to a His_6_-tag at the N-terminus.

### 2.4. Protein expression and purification

The constructs [pET28c/*Bdi*USPPase] and [pET28c/*Bdi*GolSase1] were used to transform *E. coli* BL21 Star (DE3) cells (Invitrogen). To produce the recombinant proteins, we inoculated 1 L of LB medium (supplemented with 50 µg ml^-1^ kanamycin) with a 1/100 dilution of an overnight culture. Cells were grown at 37°C and 180 rpm in an orbital shaker until OD_600nm_ was ∼0.6. Gene expression was induced with 0.5 mM isopropyl-β-D-1-thiogalactopyranoside at 25°C overnight. The cells were harvested by centrifugation at 5,000 x *g* and room temperature for 15 min and kept at -20°C until use.

The cell paste was resuspended with 25 ml of *Buffer A* [25 mM Tris-HCl pH 8.0, 300 mM NaCl, 5% (v/v) glycerol, 10 mM imidazole] supplemented with 1 mM phenylmethylsulfonyl fluoride and disrupted by sonication. The resulting suspension was centrifuged twice at 20,000 x *g* at 4°C for 10 min. The supernatant was loaded onto a 1-ml HisTrap column connected to an ÄKTA Explorer 100 purification system (GE Healthcare), previously equilibrated with *Buffer A*. The column was washed with 10 ml of *Buffer A* and the recombinant protein eluted with a linear gradient of imidazole (10 to 300 mM, 50 ml). The fractions containing the enzyme of interest were collected and concentrated. The *Bdi*GolSase1 pool was loaded onto a HiLoad 16/600 Superdex 200 pg column (GE Healthcare), previously equilibrated with *Buffer G* (50 mM HEPES-NaOH pH 8.0, 100 mM NaCl, 0.1 mM EDTA). Fractions containing GolSase activity were pooled and concentrated. Both protein preparations were supplemented with 5% (v/v) glycerol, aliquoted, and stored at -80°C until use. Under these conditions, both enzymes remained stable and active for at least 6 months.

### 2.5. Protein methods

Polyacrylamide gel electrophoresis was carried out under denaturing conditions (SDS-PAGE), as described by Laemmli (1970). Protein concentration was determined following the procedure described by Bradford (1976), using BSA as the standard.

### 2.6. Native molecular mass determination

The native molecular mass of *Bdi*USPPase and *Bdi*GolSase1 were determined by using a Superdex 200 10/300 column (GE Healthcare), equilibrated with *Buffer G*. A calibration curve was constructed by plotting the *K*_av_ values versus log_10_ of the molecular mass of standard proteins (ribonuclease, 13.7 kDa; carbonic anhydrase, 29 kDa; ovalbumin, 43 kDa; conalbumin, 75 kDa; aldolase, 158 kDa; ferritin, 440 kDa; and thyroglobulin, 669 kDa). The *K*_av_ was calculated as (V_e_-V_0_)/(V_t_-V_0_), where V_e_ is the elution volume of the protein, V_0_ is the elution volume of Dextran Blue, and V_t_ is the total volume of the column.

### 2.7. Enzyme activity assay and determination of kinetic constants

*Bdi*USPPase activity was determined in the direction of UDP-sugar synthesis, following Pi formation after hydrolysis of PPi by inorganic pyrophosphatase, using a highly sensitive colorimetric method (Fusari *et al*., 2006). The standard reaction media contained 50 mM MOPS-NaOH pH 8.0, 10 mM MgCl_2_, 1 mM UTP, 2 mM Gal1P, 0.2 mg ml^-1^ BSA, 0.025 U yeast inorganic pyrophosphatase (EC 3.6.1.1) and a proper dilution of the enzyme, in a final volume of 50 μl. Reactions were incubated for 10 min at 37°C and terminated with the addition of the Malachite Green reagent. The complex formed with Pi was measured at 630 nm in a Multiskan GO microplate reader (Thermo Scientific).

The activity of *Bdi*GolSase1 was determined in the direction of galactinol synthesis using a kinetic method coupled to pyruvate kinase and lactate dehydrogenase (Wayllace *et al*., 2012). This procedure follows the generation of UDP by measuring NADH disappearance at 340 nm. The standard assay mixture contained 50 mM HEPES-NaOH pH 7.0, 10 mM MgCl_2_, 0.5 mM PEP, 0.3 mM NADH, 1 mM UDP-Gal, 20 mM *myo-*inositol, 1 U pyruvate kinase (EC 2.7.1.40), 1 U lactate dehydrogenase (EC 1.1.1.27), 0.2 mg ml^-1^ BSA and enzyme in an appropriate dilution. Reactions were carried out for 10 min in a final volume of 50 μl at 30°C in a 384-microplate reader (Multiskan GO, Thermo Scientific).

One unit of enzyme activity (U) is defined as the amount of enzyme catalyzing the formation of 1 µmol of product per min under the above-described conditions.

Kinetic constants were determined by measuring enzyme activity at different concentrations of one substrate while keeping a fixed and saturating amount of the other one. Data of initial velocity (*v*) were plotted versus substrate concentration and fitted to a modified Hill equation: *v* = *V*_max_ [S]^*n*H^ / *S*_0.5_^*n*H^ + [S]^*n*H^, where *S*_0.5_ is the concentration of substrate (S) producing 50% of the maximal velocity (*V*_max_) and *n*_H_ is the Hill coefficient. Catalytic efficiencies were calculated as *k*_cat_ / *S*_0.5_, whereas *k*_cat_ values were obtained using the *V*_max_ and the theoretical molecular mass of each enzyme. Kinetic constants were calculated by the fitting software (Origin 8.1, OriginLab Corporation), using the mean of at least two independent sets of data, which were reproducible within ± 10%.

### 2.8. 3D modeling

The 3D structure of *Bdi*GolSase1 was modeled using a threading assembly refinement server (https://zhanglab.ccmb.med.umich.edu/I-TASSER/) (Zhang, 2008; Yang *et al*., 2011; Yang and Zhang, 2015). Based on the I-TASSER predictions, five different models of the *Bdi*GolSase1 structure were generated, and one model was finally selected based on the overall quality. Then, we used the Verify 3D server (https://servicesn.mbi.ucla.edu/Verify3D/) (Eisenberg *et al*., 1997) to test the quality of the model. Also, ligand-binding sites were predicted by the 3DLigandSite web server (http://www.sbg.bio.ic.ac.uk/3dligandsite/) (Wass *et al*., 2010).

### 2.9. Redox assays

To analyze the effect of diamide, H_2_O_2_, dithiothreitol (DTT), oxidized and reduced glutathione (GSSG and GSH, respectively) on the activity of recombinant *Bdi*GolSase1, the enzyme was incubated with increasing concentrations of the oxidizing or reducing compounds for 15 minutes at 30°C with 50 mM HEPES-NaOH pH 7.0. To remove the excess of oxidizing compounds, oxidized *Bdi*GolSase1 was ultrafiltrated by centrifuging at 10,000 x *g* and 4°C using a Vivaspin 500 device (Sartorius). To evaluate the reversibility of the process, oxidized *Bdi*GolSase1 was incubated with DTT, GSH, or *E. coli* thioredoxin (*Eco*Trx). *Eco*Trx was expressed and purified as previously described (Hartman *et al*., 2014). Reactions were incubated for 15 min at 30°C with 50 mM HEPES-NaOH pH 7.0. Aliquots were taken at regular time intervals and assayed for activity under standard conditions (see above).

## 3. Results

### 3.1. Expression, purification and characterization of Bdi*USPPase*

The sequence coding for *Bdi*USPPase was successfully cloned, and the recombinant enzyme was produced with a His_6_-tag at the N-terminus. *Bdi*USPPase has 621 amino acids and a theoretical molecular mass of 68.17 kDa. Analysis by SDS-PAGE of the purified protein showed a band of approximately 70 kDa (Fig. 1A). Gel filtration chromatography on Superdex 200 indicated *Bdi*USPPase is a monomer with a molecular mass of 86 kDa (Fig. 2B). These data are in good agreement with those reported for other plants USPPase (Supplemental Table S2).

**Figure 1.**
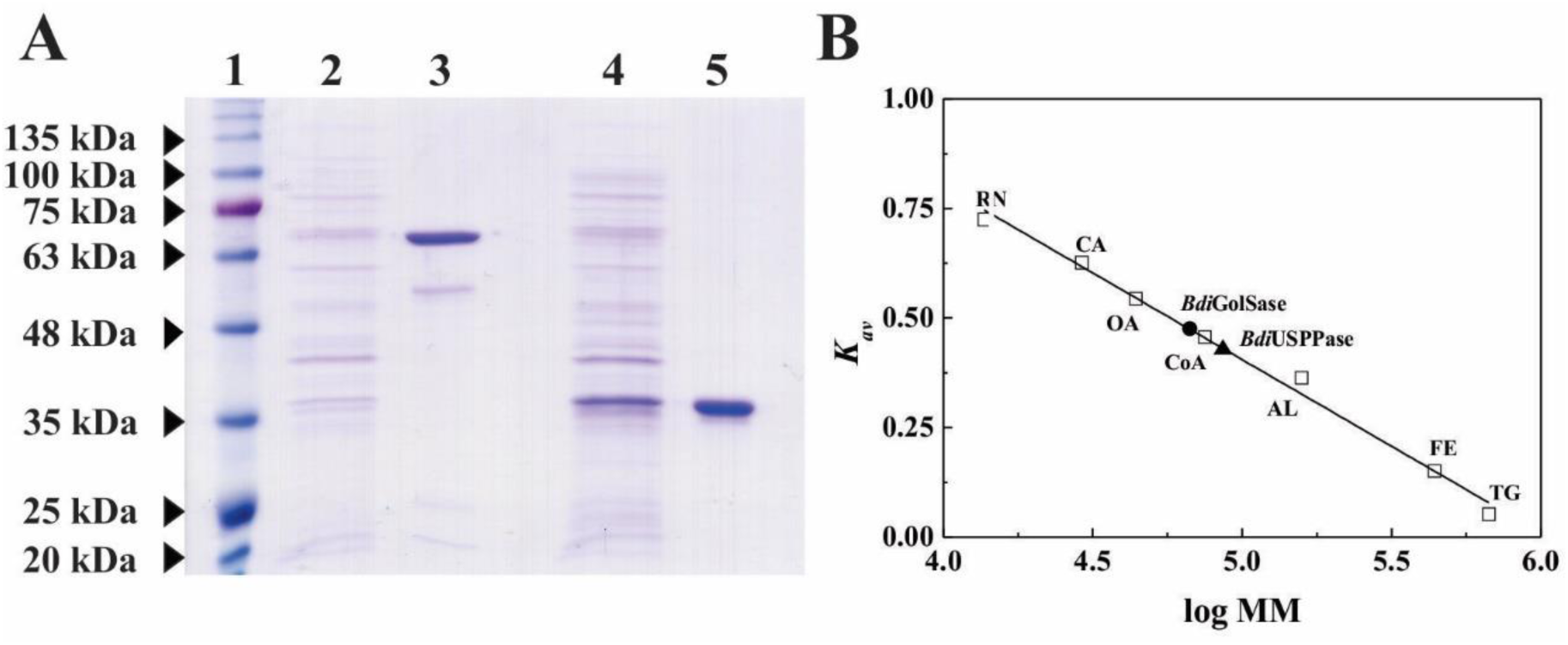
Analysis of *Bdi*USPPase and *Bdi*GolSase1 by SDS-PAGE and gel filtration chromatography. **A**. Reducing SDS-PAGE. Lane 1, molecular mass markers; lane 2, soluble fraction of recombinant cells expressing *Bdi*USPPase (crude extract); lane 3, *Bdi*USPPase purified by IMAC; lane 4, soluble fraction of recombinant cells expressing *Bdi*GolSase1 (crude extract); lane 5, *Bdi*GolSase purified by IMAC and size-exclusion chromatography. **B**. Gel filtration chromatography. Plot of *K*_av_ versus log (molecular mass). (□) Standard proteins: RN (ribonuclease, 13.7 kDa), CA (carbonic anhydrase, 29 kDa), OA (ovalbumin, 44 kDa), CoA (conalbumin, 75 kDa), AL (aldolase, 158 kDa), FE (ferritin, 440 kDa), and TG (thyroglobulin, 669 kDa); (▲) *Bdi*USPPase; (●) *Bdi*GolSase1.

**Figure 2.**
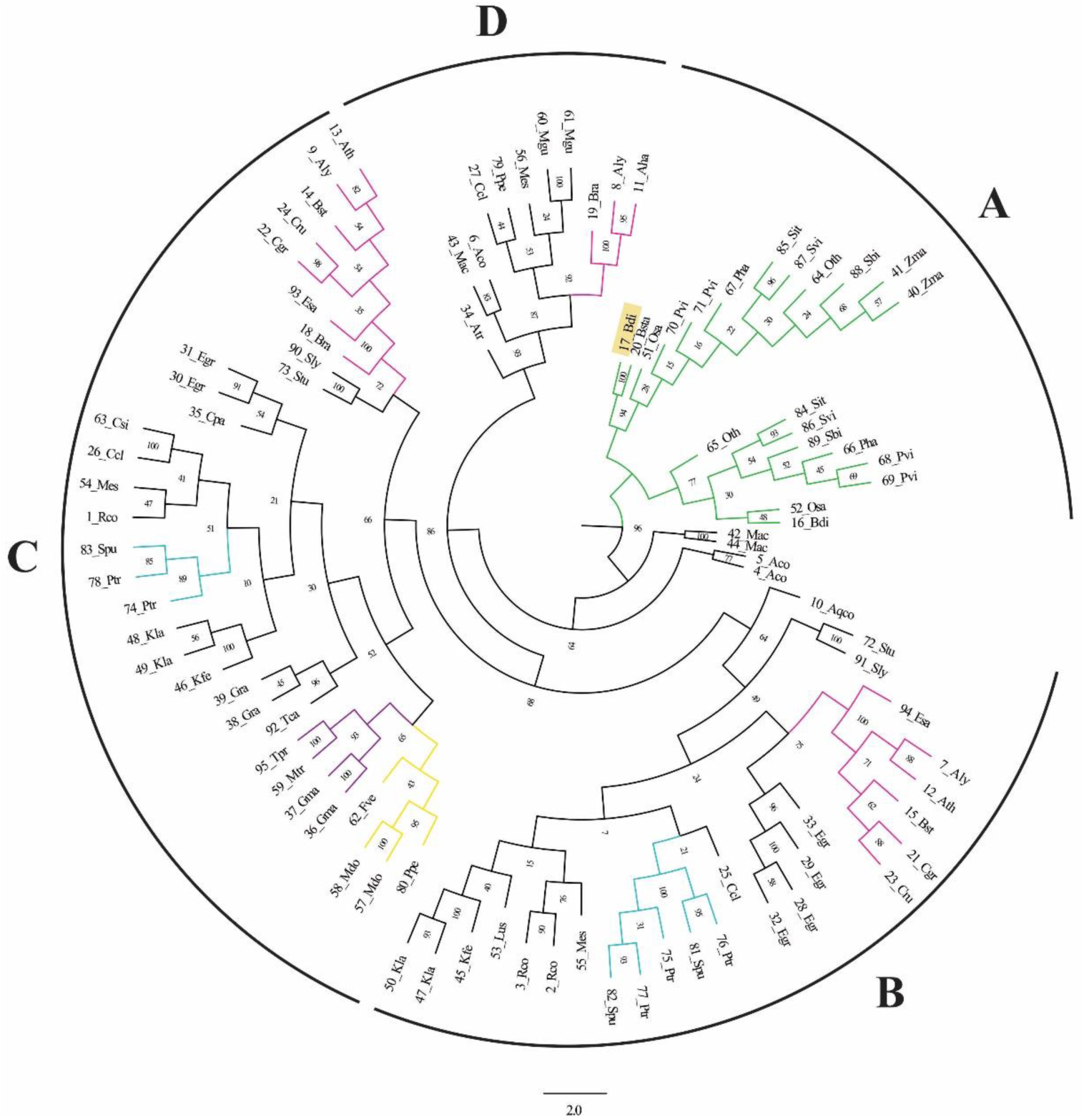
Phylogenetic analysis of GolSases and glycogenin-like proteins from plants. The tree was reconstructed using the neighbor-joining method with a bootstrap of 1,000 in Seaview 5.0.1. The sequences were number-coded for clarity (see Supplementary Table S1 for more details). *Bdi*GolSase1 is highlighted in yellow. Group A, GolSases from monocots; group B, GolSases from dicots; group C, glycogenin-like proteins from dicots; group D: GolSases from dicots. Node numbers represent the bootstrap values obtained during tree reconstruction. Families were colored as follows: Poaceae (green), Brassicaceae (pink), Rosaceae (yellow), Fabaceae (purple), and Salicaceae (light blue).

Plant USPPases show activity with different sugar 1-phosphates (Decker and Kleczkowski, 2019). To test which metabolites could be used by *Bdi*USPPase, we measured activity with a fixed concentration (2 mM) of Gal1P, Glc1P, GlcA1P, glucosamine 1-phosphate (GlcN1P), N-acetyl-glucosamine 1-phosphate (GlcNAc1P), and mannose 1-phosphate (Man1P). The highest activity was observed with Gal1P, closely followed by Glc1P (26% lower), while that with other sugar-1P (GlcA1P, GlcN1P, GlcNAc1P, and Man1P) was 5-to 17-fold lower than with Gal1P (Supplemental Figure S1). Then, we determined the kinetic parameters for *Bdi*USPPase in the direction of UDP-sugar synthesis. We could calculate *V*_max_ and *S*_0.5_ values when using Gal1P, Glc1P, and GlcA1P as substrates (Table 1) but not for GlcN1P, GlcNAc1P, and Man1P because curves did not reach saturation (Supplemental Figure S2). The *V*_max_ of *Bdi*USPPase was highest with Gal1P, slightly lower with Glc1P and 5-fold lower with GlcA1P. The apparent affinities (estimated from *S*_0.5_ values) for Gal1P and Glc1P were similar but 5-fold lower for GlcA1P. The catalytic efficiency of *Bdi*USPPase was almost the same for Gal1P and Glc1P, but one order of magnitude lower for GlcA1P (Table 1). Overall, these results are similar to those obtained for USPPases from other plant species (Supplemental Table S2) and indicate *Bdi*USPPase could provide one of the substrates (i.e., UDP-Gal) for galactinol synthesis.

**Table 1.** Kinetic parameters for the substrates of *Bdi*USPPase. Reactions were performed in the direction of UDP-sugar synthesis. Kinetic constants were calculated by fitting experimental data to a modified Hill equation, as described in section 2.7. Parameters were calculated using the mean of at least two independent datasets ± standard error.

| Substrate | $S_{0.5}$<br>(mM) | $n_H$ | $V_{max}$<br>(U mg<sup>-1</sup>) | Catalytic efficiency<br>(M<sup>-1</sup> s<sup>-1</sup>) |
| --- | --- | --- | --- | --- |
| Gal1P | 0.25 ± 0.03 | 1.0 ± 0.1 | 1024 ± 95 | 4.7 x 10<sup>6</sup> |
| UTP<sub>Gal1P</sub> | 0.109 ± 0.007 | 1.4 ± 0.1 | | 1.1 x 10<sup>7</sup> |
| Glc1P | 0.28 ± 0.02 | 1.4 ± 0.1 | 757 ± 27 | 3.1 x 10<sup>6</sup> |
| UTP<sub>Glc1P</sub> | 0.11 ± 0.01 | 1.9 ± 0.2 | | 7.8 x 10<sup>6</sup> |
| GlcA1P | 1.4 ± 0.3 | 1.2 ± 0.2 | 183 ± 12 | 1.5 x 10<sup>5</sup> |
| UTP<sub>GlcA1P</sub> | 0.14 ± 0.01 | 1.4 ± 0.2 | | 1.5 x 10<sup>6</sup> |

### 3.2. *Phylogenetic analysis of* Bdi*GolSase1*

To better understand the function and properties of *Bdi*GolSase1, we first reconstructed a phylogenetic tree with GolSase and glycogenin-like protein sequences from different plant species. The resulting tree was divided into four main groups, namely A to D (Fig. 2). GolSase sequences from monocots (Poaceae family) clustered in a single clade (A), as observed in previous studies (dos Santos *et al*., 2011; Sengupta *et al*., 2012; Zhou *et al*., 2014). The sequences from dicots were further separated into three clades (B to D). Groups B and D contain GolSase sequences, while group C includes sequences annotated as glycogenin-like proteins (Fig. 2). The two sequences annotated as GolSases in the *B. distachyon* genome (16_Bdi and 17_Bdi, Supplemental Table S1) were located in the clade containing monocot sequences (A), although in different subgroups (Fig. 2).

### 3.3. *Expression, purification, and characterization of* Bdi*GolSase1*

The *BdiGolSase1* gene encodes a protein of 337 amino acids (17_Bdi; Supplemental Table S1 and Fig. 2), which has a theoretical molecular mass of 37.9 kDa. Recombinant *Bdi*GolSase1 was highly purified by IMAC and gel filtration chromatography, as assessed by SDS-PAGE (Fig. 1A). The enzyme eluted from the analytical gel filtration chromatography on Superdex 200 as a monomer of 45 kDa (Fig. 2B), which agrees with results obtained for GolSases from other plant species (Supplemental Table S3).

Then, we analyzed the effect of divalent cations on *Bdi*GolSase1. The enzyme exhibited a basal activity of 14.9 U mg^-1^ when assayed without metal ions. Such a value increased to 28.6 U mg^-1^ with Mg^2+^ and 41.4 U mg^-1^ with Mn^2+^ (10 mM each). The apparent affinity for Mn^2+^ was 10-fold higher than for Mg^2+^ (*A*_0.5_ values of 0.09 and 1.4 mM, respectively; Supplemental Fig. S3). Since the cytosolic concentration of Mn^2+^ in plant cells is in the low (0.3-0.8) µM range (Quiquampoix *et al*., 1993; Pittman, 2005), and Mg^2+^ concentrations vary from 15 to 25 mM (Moomaw and Maguire, 2008; Waters, 2011), we continued the kinetic characterization of *Bdi*GolSase1 with 10 mM Mg^2+^. Afterward, we measured *Bdi*GolSase1 activity at different pH and temperature values. As shown in Supplemental Fig. S4, the activity was optimal at pH values ranging from 7.0 to 9.0 and at 35°C. Data of *Bdi*GolSase1 activity at different temperatures were used to calculate the enzymes’s activation energy (Segel *et al*., 1993), which resulted in 31.76 kJ mol^-1^ (Supplemental Fig. S4A, inset).

Table 2 shows the kinetic parameters obtained for *Bdi*GolSase1, both with UDP-Gal (the natural substrate) and UDP-glucose (UDP-Glc, used to test the specificity of the recombinant protein). When using UDP-Gal as a substrate, the enzyme exhibited a 40-fold lower *S*_0.5_ and reached a 26-fold higher *V*_max_ than with UDP-Glc. Similarly, the apparent affinity for *myo*-inositol was 8-fold higher with UDP-Gal than with UDP-Glc. The catalytic efficiency was three orders of magnitude higher when using UDP-Gal compared with that obtained with UDP-Glc. No activity was detected when using UDP-GlcA or ADP-Glc as substrates (data not shown). We also tested the effect of several hexose-phosphates (glucose 6-phosphate, mannose 6-phosphate, fructose 6-phosphate, fructose 1,6-bisphosphate, glucosamine 6-phosphate) on the activity of *Bdi*GolSase1, measured with UDP-Gal and *myo*-inositol. However, no significant differences were observed when they were present into the assay mixture (5 mM each, data not shown).

**Table 2.** Kinetic parameters for the substrates of *Bdi*GolSase1. Reactions were performed in the direction of galactinol synthesis. Kinetic constants were calculated by fitting experimental data to a modified Hill equation, as described in section 2.7. Parameters were calculated using the mean of at least two independent datasets ± standard error.

| Substrate | $S_{0.5}$<br>(mM) | $n_H$ | $V_{\max}$<br>(U mg<sup>-1</sup>) | Catalytic efficiency<br>(M<sup>-1</sup> s<sup>-1</sup>) |
| --- | --- | --- | --- | --- |
| UDP-Gal | 0.06 ± 0.01 | 1.5 ± 0.3 | 28.6 ± 0.9 | 3.2 x 10<sup>5</sup> |
| *myo*-inositol UDP-Gal | 2.5 ± 0.3 | 1.1 ± 0.2 | | 7.6 x 10<sup>3</sup> |
| UDP-Glc | 2.6 ± 0.5 | 1.1 ± 0.1 | 1.1 ± 0.2 | 2.8 x 10<sup>2</sup> |
| *myo*-inositol UDP-Glc | 20 ± 3 | 1.1 ± 0.1 | | 3.7 x 10<sup>1</sup> |

### 3.4. *3D modeling of* Bdi*GolSase1*

To build a 3D model of *Bdi*GolSase1, we first searched for solved structures (templates) in the Protein Data Bank using the BLAST algorithm from the NCBI server. This search resulted in the crystal structures of human (4UEG) and rabbit (1LL0) glycogenins having the best fit. The sequence of *Bdi*GolSase1 has a relatively low identity (24-28%) with these enzymes, which prevents the use of homology modeling. Therefore, we used protein threading (fold recognition) with the Iterative Threading ASSEmbly Refinement (I-TASSER) on-line server to construct a model for *Bdi*GolSase1. The output of this server included five full-length atomic models ranked based on cluster density, from which we selected model 1 to continue with the structural analysis. The *Bdi*GolSase1 3D model was obtained in complex with UDP-Glc and Mn^2+^, using the resolved crystal structures of rabbit and human glycogenins as threading templates. The structural model displays ten α-helices, eight β-strands, and a large number of loose coils. The N-terminal portion of *Bdi*GolSase1 shows a Rossmann-fold composed of five β-strands, with the last strand in antiparallel direction (Fig. 3A).

**Figure 3.**
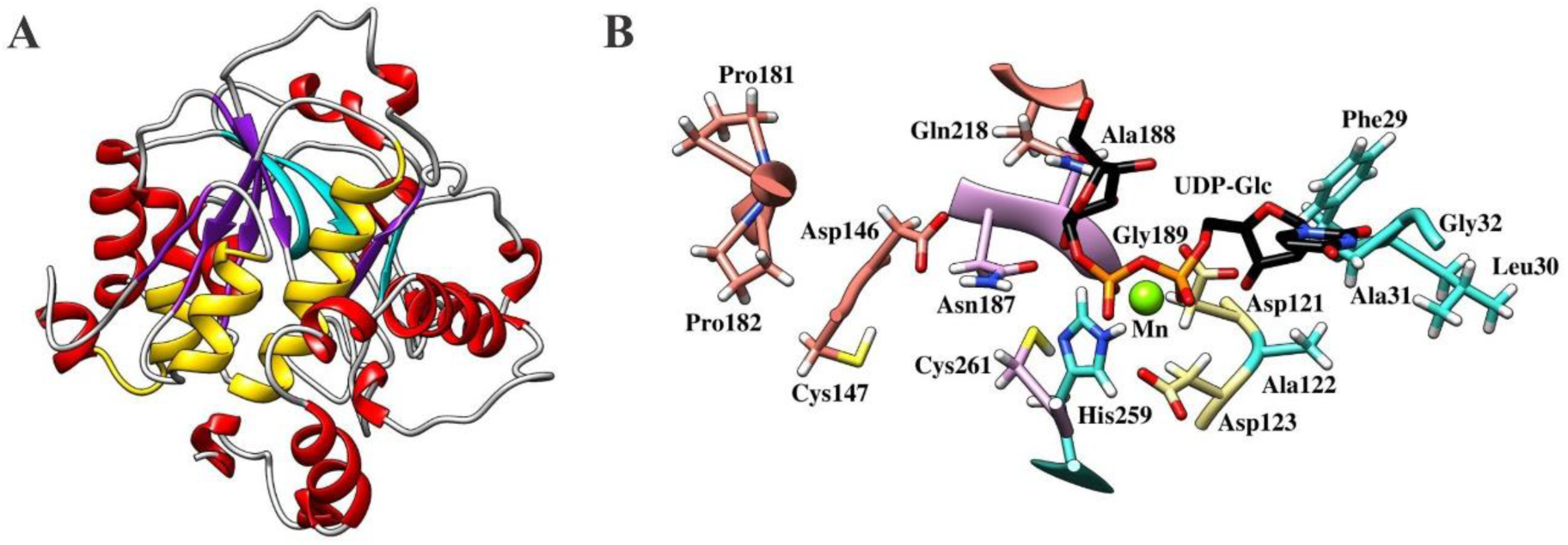
The structural model of *Bdi*GolSase1 obtained by protein threading (fold recognition). **A**. The picture shows the Rossmann fold within the N-terminal domain of *Bdi*GolSase1, with β-strands highlighted in purple and α-helices in yellow. **B**. Close up view of the model complexed with UDP-Glc (colored by atom type) and Mn^2+^ (green ball). The putative catalytic site’s residues were colored as follows: yellow, residues that bind the metal ion; light blue, residues that interact with the UDP-sugar; pink, residues that bind *myo*-inositol; violet, residues that bind both substrates.

A close inspection of the protein sequence showed high conservation of motifs found in enzymes from the GT8 family, including the DxD, HxxGxxKPW, GLG, NAG, and FLAG domains (Supplemental File S1), also described in previous publications (Sengupta *et al*., 2012; Salvi *et al*., 2016; Fan *et al*., 2017). Most of these regions are buried inside the protein, forming the putative catalytic pocket (Fig. 3B). The DxD motif (Asp^121^-Ala^122^-Asp^123^) is located between strands β4 and β5 of the Rossman-fold and is part of the catalytic domain (Fig. 3B). The two Asp residues interact with the Mn^2+^ ion, while Ala^122^ binds to the ribose moiety of UDP-Glc. Other amino acids that might be involved in the binding of UDP-Glc are Phe^29^, Leu^30^, Ala^31^, Gly^32^, Asn^187^, Ala^188^, Gly^189^, His^259^, and Cys^261^ (Fig. 3B). The I-TASSER server does not provide information about the *myo*-inositol binding site since there are no resolved crystal structures for glycosyltransferases using this substrate. However, submission of the model obtained for *Bdi*GolSase1 to the 3DLigandSite server allowed us to identify putative candidates involved in *myo*-inositol binding: Asp^146^, Cys^147^, Pro^181^, Pro^182^, Asn^187^, Ala^188^, Gly^189^, Gln^218^ and Cys^261^ (Fig. 3B).

### 3.5. *Redox regulation of* Bdi*GolSase1*

We identified six Cys residues in the *Bdi*GolSase1 3D model (Supplemental Fig. S5), being Cys^261^ located in the putative catalytic site (Fig. 3B). Therefore, we tested the effect of oxidizing and reducing agents on the activity of the recombinant enzyme. *Bdi*GolSase1 was incubated with increasing concentrations of different oxidizing agents, as described in Section 2.9. Incubation of the recombinant enzyme with 1 mM diamide or 0.5 mM H_2_O_2_ produced 40% inhibition, while the activity was negligible after incubation with 10 mM of both oxidants (Supplemental Fig. S6). Incubation with 0.1 mM GSSG showed 40% inhibition of *Bdi*GolSase1 activity, while higher concentrations produced the same effect. Exposure to the reducing agents DTT and GSH slightly increased (∼25%) the activity of the recombinant enzyme (Supplemental Fig. S6).

After that, we tested if the activity of oxidized *Bdi*GolSase1 could be recovered by incubation with chemical or biological reducing agents. The enzyme was first treated with 1 mM diamide or H_2_O_2_ and then incubated with 1 mM DTT or 50 µM *Eco*Trx. We observed that activity of the diamide-oxidized *Bdi*GolSase1 was recovered by both DTT and *Eco*Trx (Fig. 4). Conversely, the H_2_O_2_-oxidized *Bdi*GolSase1 could not be rescued from inhibition by any of the tested reducing agents, suggesting an irreversible, deleterious process caused by the oxidant. The GSSG-treated enzyme showed no significant variations after incubation with GSH or *Eco*Trx (Supplemental Fig. S7).

**Figure 4.**
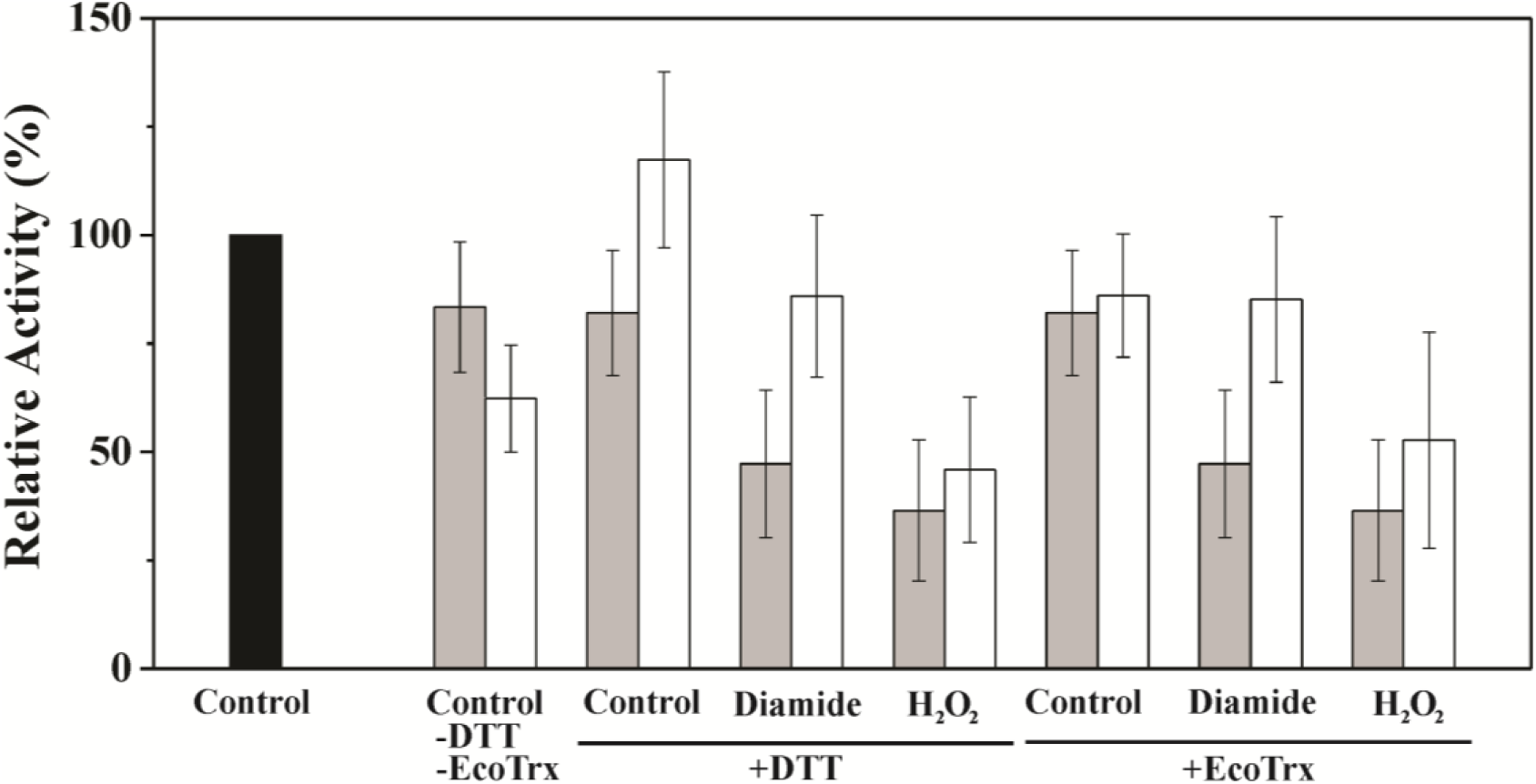
Recovery of oxidized *Bdi*GolSase1 by reduction with DTT or *Eco*Trx. Enzyme activity data were related to the value obtained at the beginning of each assay. Black bars, relative activity at the beginning of the assay; gray bars, relative activity after incubation with the oxidizing agents (diamide or H_2_O_2_); white bars, relative activity of the oxidized samples after incubation with the reducing agents (DTT or *Eco*Trx). Data are the mean of two independent data sets ± standard error.

## 4. Discussion

The number of studies on enzymes involved in Raf metabolism is relatively scarce (Ende, 2013; Elsayed *et al*., 2014; Sengupta *et al*., 2015). Herein, we report the recombinant production of two enzymes from *B. distachyon* involved in Raf synthesis, *Bdi*USPPase, and *Bdi*GolSase1. The molecular mass of *Bdi*USPPase determined by SDS-PAGE and gel filtration chromatography (Fig. 1) is in good agreement with the theoretical value and with data reported for USPPases from dicot species (Kotake *et al*., 2004; Dai, 2006; Litterer *et al*., 2006*b,a*). Recombinant *Bdi*USPPase is active as a monomer and, unlike parasite and plant UDP-Glc pyrophosphorylases, there is no evidence for oligomerization (Damerow *et al*., 2010; Decker and Kleczkowski, 2019). The reaction catalyzed by USPPase is reversible (Kleczkowski *et al*., 2010, 2011) and Mg^2+^-dependent (Litterer *et al*., 2006*a*; Mu *et al*., 2009; Yang *et al*., 2010), following an ordered bi-bi mechanism (Kleczkowski *et al*., 2011). In this work, we kinetically characterized *Bdi*USPPase in the direction of UDP-sugar synthesis (Table 1). The enzyme has a similar apparent affinity for Gal1P and Glc1P, but 5-fold lower for GlcA1P. These parameters are consistent with those previously reported for USPPases from other plants (Supplemental Table S2). Interestingly, the *S*_0.5_ for UTP did not significantly change, regardless of the sugar-1P used (Table 1). Kinetic parameters could not be determined for GlcN1P, GlcNAc1P, and Man1P, because curves did not reach saturation at the highest concentration tested, indicating the apparent affinity for these substrates is considerably lower than for Gal1P and Glc1P.

The *B. distachyon* genome has two genes encoding putative GolSases, sharing ∼74% amino acid identity (Supplemental File S1). We reconstructed a phylogenetic tree using 95 GolSase and glycogenin-like protein sequences from several plant species. Sequences were divided into four main groups, and both *B. distachyon* GolSases were located in the clade containing monocot GolSases (group A), although in different subgroups (Fig. 2). The presence of various *GolSase* genes in the same species suggests gene duplication, which leads to molecular innovations in higher organisms (Blanc and Wolfe, 2004; Ganko *et al*., 2007). The subfunctionalization of GolSases (Rastogi and Liberles, 2005), not only in *B. distachyon* but also in other plant species (Fig. 2), might allow plants tolerate different abiotic stress conditions (Fan *et al*., 2017). As previously mentioned, different GolSase genes are expressed in *A. thaliana* plants exposed to drought, heat, and high salinity (*AtGolS1* and *AtGolS2*) or cold (*AtGolS3*; Taji *et al*., 2002).

Analysis by SDS-PAGE and gel filtration chromatography (Fig. 1) showed that *Bdi*GolSase1 is a monomer, which is consistent with data previously reported for GolSases from other plant species (Smith *et al*., 1991; Liu *et al*., 1995; Zhou *et al*., 2017; Kito *et al*., 2018). The activity of *Bdi*GolSase1 is higher in the presence of Mn^2+^ than Mg^2+^ (Supplemental Fig. S3), being the former required for optimal catalysis (Handley and Pharr, 1982; Webb, 1983; Smith *et al*., 1991; Liu *et al*., 1995; Li *et al*., 2011). The optimal pH and temperature values determined for *Bdi*GolSase1 (Supplemental Fig. S4) are similar to those reported for other plant GolSases (Bachmann *et al*., 1994; Liu *et al*., 1995; Ribeiro *et al*., 2000; Wakiuchi *et al*., 2003). The activation energy calculated for *Bdi*GolSase1 is similar to that obtained for aldose-6-phosphate reductase from apple leaves (Figueroa and Iglesias, 2010). The kinetic characterization of *Bdi*GolSase1 showed *S*_0.5_ and *V*_max_ values that resemble those described for GolSases from dicot species; remarkably, the *S*_0.5_ of *Bdi*GolSase1 for UDP-Gal is the lowest reported so far (Table 2 and Supplemental Table S3).

The protein sequence of *Bdi*GolSase1 showed well-conserved motifs present in the GT8 family (Sengupta *et al*., 2012). This information, combined with the structural model obtained by protein threading (Fig. 3), allowed us to determine the residues putatively involved in substrate and metal binding. It has been shown that the two Asp residues from the DxD motif present in glycosyltransferases, as well as both phosphate groups from the NDP-sugar, are involved in the coordination of the divalent metal ion (Breton and Imberty, 1999; Ünligil and Rini, 2000; Breton *et al*., 2006). This is consistent with our modeling results for *Bdi*GolSase1, where we observed that Asp^121^ and Asp^123^ bind to the Mn^2+^ ion (Fig. 3B).

A common mechanism affecting enzyme activity *in vivo* is the oxidation of Cys residues, mainly caused by reactive oxygen species, reactive nitrogen species, and GSSG (Piattoni *et al*., 2013). *Bdi*GolSase1 contains six Cys residues (Supplemental Fig. S5); one of them (Cys^261^) is part of the putative catalytic site and might be involved in the binding of both substrates, UDP-Gal and *myo*-inositol (Fig. 3B). Interestingly, the recombinant enzyme was inactivated by oxidation with diamide, and the activity was recovered by DTT and *Eco*Trx (Fig. 4), suggesting the existence of a redox regulatory mechanism. Unlike diamide, which is considered a mild-oxidant, H_2_O_2_ is a strong oxidant (Burek *et al*., 2019). Incubation of the recombinant enzyme with H_2_O_2_ caused irreversible inactivation (Fig. 4), probably due to thiol oxidation to sulfinic or sulfonic acids, oxidation states that cannot be reversed by DTT or Trx (Kiley and Storz, 2004; Poole *et al*., 2004). While oxidative modification of one or a few surface-exposed amino acids generally has little effect on the stability of the enzyme, oxidation becomes more critical if the modified amino acid lies within the active site or is actively involved in the catalytic mechanism (Burek *et al*., 2019).

To conclude, in this work, we biochemically characterized two enzymes involved in Raf metabolism, *Bdi*USPPase, and *Bdi*GolSase1. To the best of our knowledge, this is the first comprehensive kinetic characterization of a GolSase from a monocot species. The combination of structural and biochemical data allowed us to establish structure-to-function relationships in *Bdi*GolSase. Overall, our work lays the ground to better understand the synthesis of galactinol and Raf in plants.

## Contributions

AAI and CMF conceived the study; RIM and MPM performed experiments; all authors analyzed data; RIM, AAI, and CMF wrote the manuscript; all authors approved the final version.

## Acknowledgments

This work was supported by grants from ANPCyT (PICT-2017-1515 and PICT-2018-00929 to AAI and PICT-2018-00865 to CMF) and UNL (CAI+D). CMF is funded by the Max Planck Society (Partner Group for Plant Biochemistry). RIM and MPM are Fellows, and AAI and CMF are Researchers from CONICET.

## Abbreviations

*Bdi*USPPase: *Brachypodium distachyon* UDP-sugar pyrophosphorylase
*Bdi*GolSase1: *Brachypodium distachyon* galactinol synthase 1
DTT: dithiothreitol
*Eco*Trx: *Escherichia coli* thioredoxin
Gal1P: galactose 1-phosphate
Glc1P: glucose 1-phosphate
GlcA1P: glucuronic acid 1-phosphate
GlcN1P: glucosamine 1-phosphate
GlcNAc1P: N-acetyl-glucosamine 1-phosphate
GSH: reduced glutathione
GSSG: oxidized glutathione
Man1P: mannose 1-phosphate
Raf: raffinose
UDP-Gal: UDP-galactose
UDP-Glc: UDP-glucose.

## Notes

### Competing Interest Statement

The authors have declared no competing interest.

