## Supplemental Figures S1-S7 for "Biochemical characterization of recombinant UDP-sugar pyrophosphorylase and galactinol synthase from *Brachypodium distachyon*"

**Supplemental Figure S1**

Activity of *Bdi*USPPase with different sugar-1P. Enzyme activity was assayed in the direction of UDP-sugar synthesis, with 1 mM UTP and 2 mM sugar-1P. Gal1P, galactose 1-phosphate; Glc1P, glucose 1-phosphate; GlcA1P, glucuronic acid 1-phosphate; Man1P, mannose 1-phosphate; GlcNAc1P, N-acetyl-glucosamine 1-phosphate; GlcN1P, glucosamine 1-phosphate. Data are the mean of two independent data sets ± standard error.

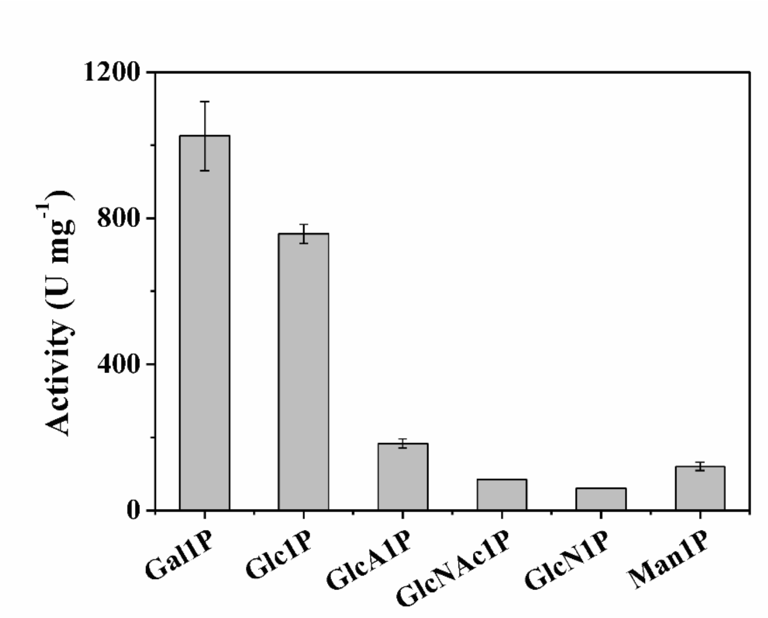

**Supplemental Figure S2**

Activity of *Bdi*USPPase with increasing concentrations of GlcNAc1P, GlcN1P, and Man1P. Enzyme activity was assayed in the direction of UDP-sugar synthesis, with 1 mM UTP and variable concentrations of the sugar-1P. Data are the mean of two independent data sets ± standard error.

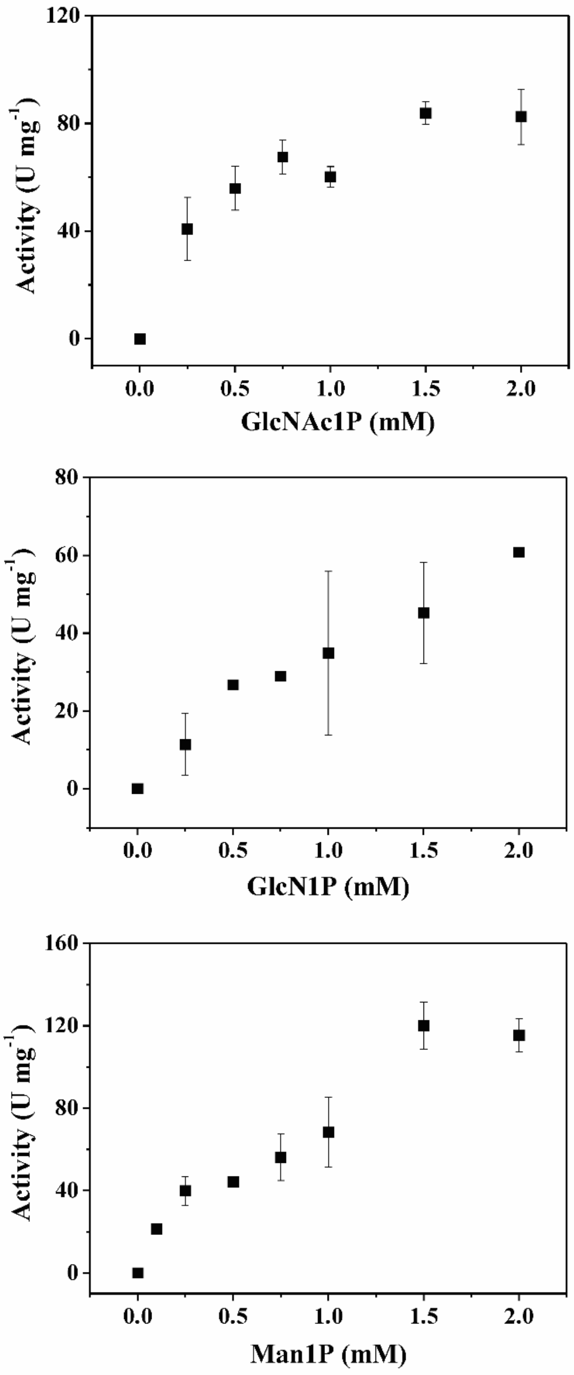

**Supplemental Figure S3**

*Bdi*GolSase activity was assayed with increasing concentrations of $Mn^{2+}$ (black squares) or $Mg^{2+}$ (white squares). Constants were calculated by fitting experimental data to a modified Hill equation, using the mean of at least two independent data sets ± standard error.

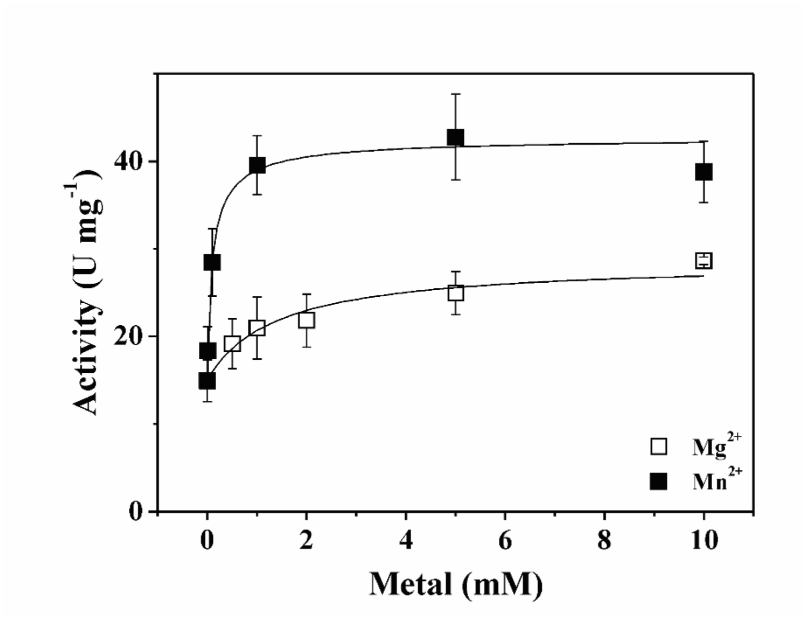

**Supplemental Figure S4**

Analysis of *Bdi*GolSase activity at different pH (A) and temperature (B) values. A, inset: Arrhenius plot for *Bdi*GolSase using data of enzyme activity between 15 and 35°C. Buffers used in (B) were: HEPES-NaOH (pH 6.8-8.2), Bis-Tris Propane-NaOH (BTP, pH 6.1-10.1), MES-NaOH (pH 5.0-6.8), CAPS-NaOH (pH 9.5-11.0) and Tris-HCl (pH 7.5-9). Data are the mean of two independent data sets ± standard error.

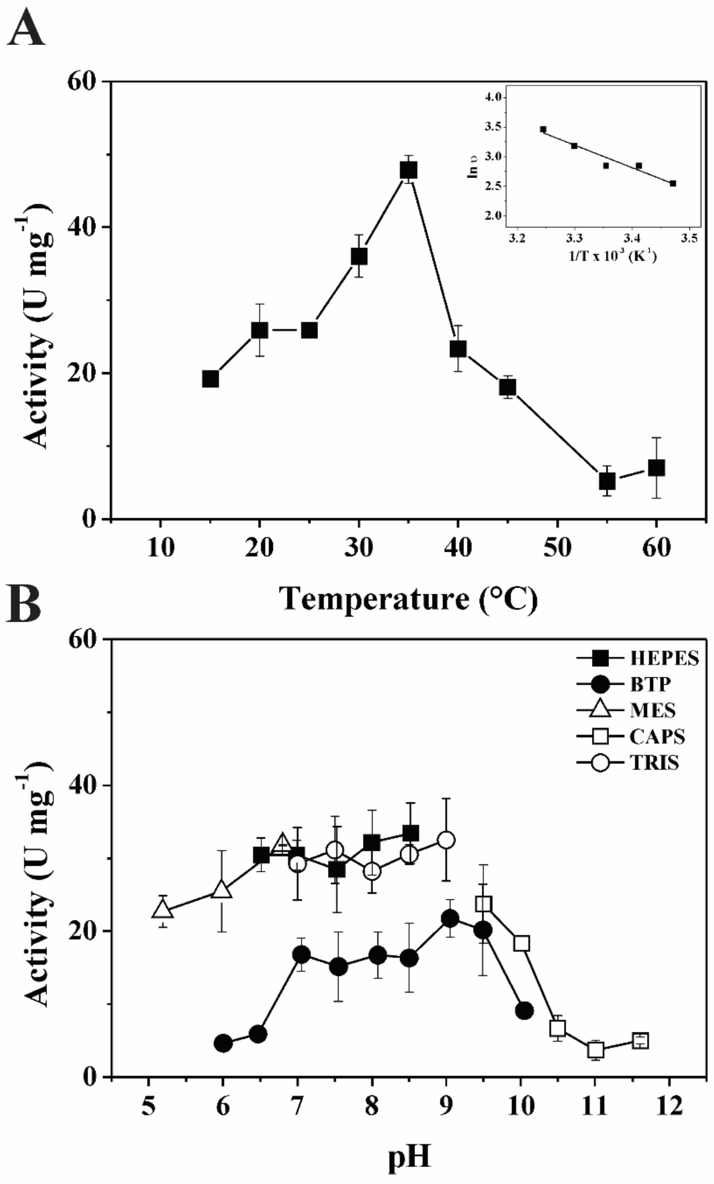

**Supplemental Figure S5**

Structural model of *Bdi*GolSase where cysteine residues are shown in different color and their positions labeled.

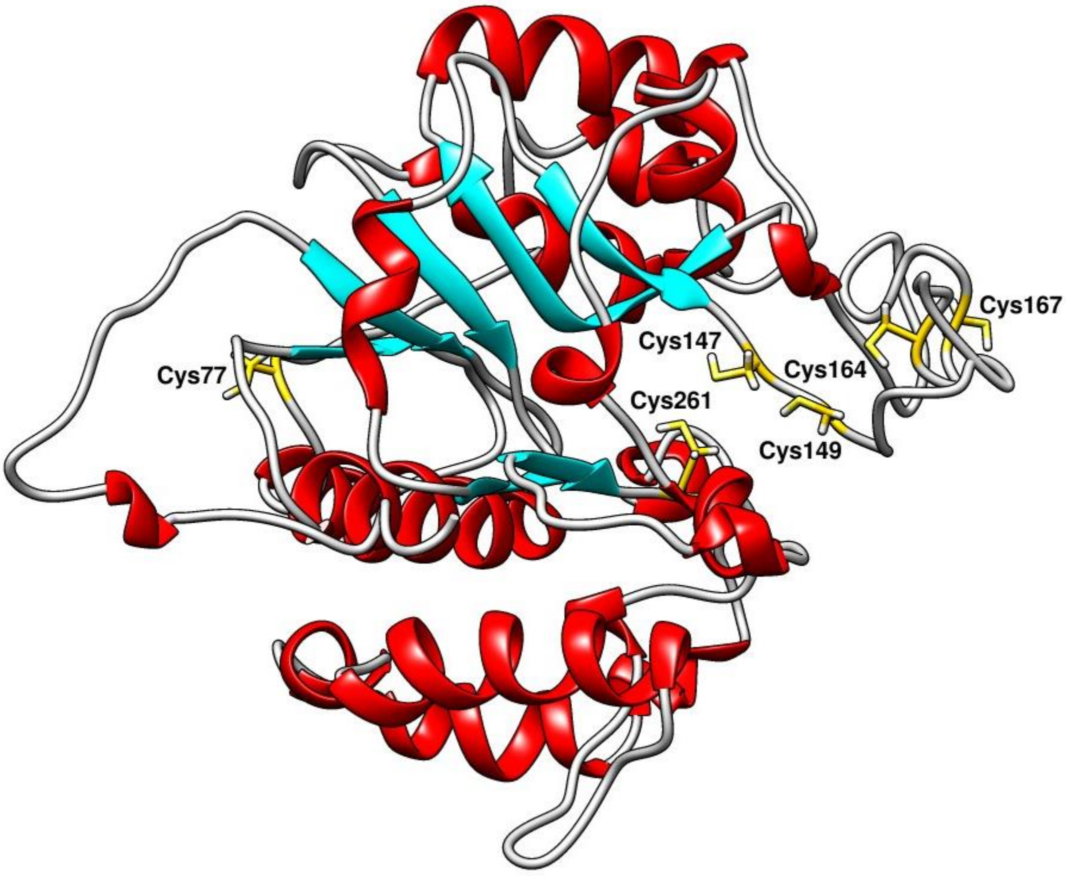

**Supplemental Figure S6**

Effect of diamide, $H_2O_2$, GSSG, DTT and GSH on the activity of *Bdi*GolSase. Data are the mean of at least two independent data sets ± standard error. *, p<0.05; **, p<0.01; ***, p<0.001 (t-test).

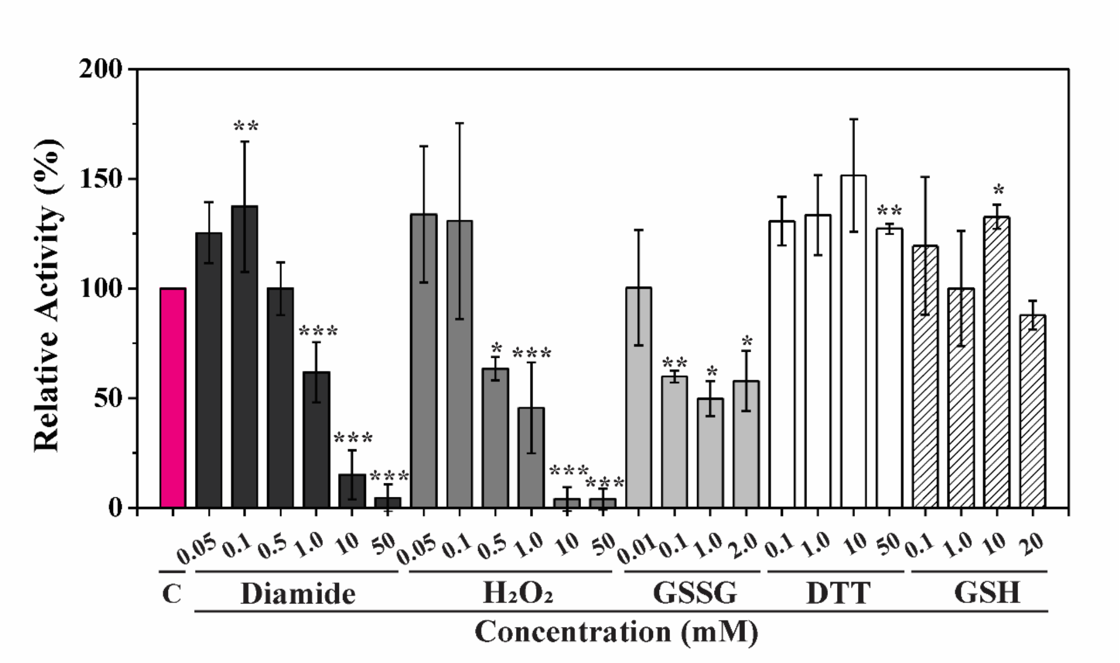

**Supplemental Figure S7**

Treatment of GSSG-oxidized *Bdi*GolSase with DTT or *Eco*Trx. Enzyme activity data were related to the value obtained at the beginning of each assay. Data are the mean of two independent data sets ± standard error.

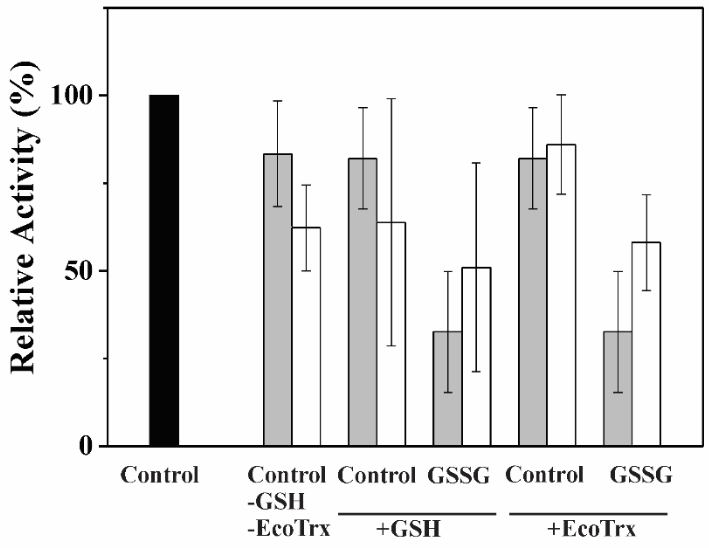
