## Supplemental File S1 for "Biochemical characterization of recombinant UDP-sugar pyrophosphorylase and galactinol synthase from *Brachypodium distachyon*"

```
                    10        20        30        40        50        60        70        80        90       100
            ....|....|....|....|....|....|....|....|....|....|....|....|....|....|....|....|....|....|....|....|
1_Rco       -----MAP-ELVHAALKPASF-TKPPTL----PSRAYVTFLAGNGDYIKGVVGLAKGLRKVKTAYPLVVAVLPDVPEEHRKILESQGCIVREIEPVYPPE
2_Rco       ----MAPHLASANLAANTNS---LVKQA--SISSCAYVTFLAGDGDYVKGVVGLAKGLRKVKSKYPLVVAILPDVPEDHRKILVSQGCIVKEIEPVYPPE
3_Rco       ----MAPHLTSANLAANTNS---LVKQA--SISSCAYVTFLAGNGDYVKGVVGLAKGLRKVNSKYPLVVAILPDVPEDHRKILVSQGCIIKEIEPVYPPE
4_Aco       ---------MAPEIVGKAAAVAKAVAKA---KAKNAYVTFLAGDGDYVKGVVGLAKGLRKVGSAYPLVVAVLPDVPESHRRLLRSQGCIVREIEPVYPPE
5_Aco       ---------MDTKIVKKLSPNSKS----------NVYVTFLAGDGDYVKGVVGLAKGLRKVGSAYPLVVAVLDDVPESHRRLLESQGCIVREIEPVHPPE
6_Aco       -----MAPIVPTAPAAGKSGGRVMKS--------RAYVTFLAGDGDYVKGVVGLAKGLRKVGSAYRLVVAVLPDVPEAHRRILATQGCVVREIEPVYPPE
7_Aly       ---------MAPEINTKLTR--PVLSATAYGGEKRAYVTFLAGTGDYVKGVVGLAKGLRKAKSKYPLVVAVLPDVPEDHRKQLVDQGCVVKEIEPVYPPE
8_Aly       -----MAP-QIPVNSIYLSEKAHQAP------PKRAYVTFLAGNGDYVKGVVGLAKGLRKVKSAYPLVVAMLPDVPEEHREILRSQGCILREIEPVYPPE
9_Aly       -----MAPGLTPTADAESTVMITKPLLPSVQDSDRAYVTFLAGNGDYVKGVVGLAKGLRKVKSAYPLVVAILPDVPEEHRRILMEQGCIVREIEPVYPPE
10_Aqco     ---------MAPNIISTATKTVGFTKQA--SIPSRAFVTFLAGNGDYVKGVVGLAKGLRKVKSAYPLVVAILPDVPEEHRRILESQGCIVREIEPVYPPE
11_Aha      -----MAP-EIPINSIYLSEKAHQAP------PRRAYVTFLAGNGDYVKGVVGLAKGLRKVKSAYPLVVAMLPDVPEEHREILRSQGCILREIEPVYPPE
12_Ath      ---------MAPEINTKLTV--PVHSAT--GGEKRAYVTFLAGTGDYVKGVVGLAKGLRKAKSKYPLVVAVLPDVPEDHRKQLVDQGCVVKEIEPVYPPE
13_Ath      -----MAPGLTQTADAMSTVTITKPSLPSVQDSDRAYVTFLAGNGDYVKGVVGLAKGLRKVKSAYPLVVAMLPDVPEEHRRILVDQGCIVREIEPVYPPE
14_Bst      -----MAPELTQNAATKSTVTITKPSPP-IQGSDRAYVTFLAGNGDYVKGVVGLAKGLRKVKSAYPLVVAILPDVPEEHRRILVEQGCIVREIEPVYPPE
15_Bst      ---------MAPENNTKLNNNGPVVSATAYGGEKRAYVTFLAGNGDYVKGVVGLAKGLRKAKSKYPLVVAVLPDVPEDHRKQLVDQGCLVKEIEPVYPPE
16_Bdi      -----MAPLMNGSCKSEEKK------------LPGAYVTFLAGTGDYWKGVVGLAKGLRAVKSAHPLVVAVLPDVPASHRQILASQGCLVRAIQPVYPPE
17_Bdi      -----MAP-ELSRKMTTAKAAAAAV-----KPATKAFVTFLAGDGDYWKGVVGLAKGLRKAGSAYPLVVAMLPDVPESHRRILASQGCILREIVPVYPPE
18_Bra      -----MAPELTQTTTVKSAVTITKPSPP-VHG-DRAYVTFLAGNGDYVKGVVGLAKGLRKVKSAYPLVVAILPDVPEEHRRVLVEQGCIVREIEPVYPPE
19_Bra      -----MAP-EISVNTLSISEKVHLAP------SKRAYVTFLAGDGDYVKGVVGLAKGLRKVKSAYPLVVAMLPDVPEEHREILRSQGCVLREIDPVHPPD
20_Bsta     -----MAP-ELSGKMTTAK-AAAAV-----KPATKAFVTFLAGDGDYWKGVVGLAKGLRKAGSAYPLVVAMLPDVPESHRRILASQGCILREIVPVYPPE
21_Cgr      ---------MAPEINTKLTT-GPVISATAYGGEKRAYVTFLAGNGDYVKGVVGLAKGLRKAESKYPLVVAVLPDVPEDHRKQLVDQGCVVKEIEPVYPPE
22_Cgr      -----MAPELTQTAAAKSTVTITKPSPPTIYGGGRAYVTFLAGNGDYVKGVVGLAKGLRKVKSAYPLVVAILPDVPEEHRRILVEQGCIVREIEPVYPPD
23_Cru      ---------MAPEINTKLTT-GPVIPATAYGGEKRAYVTFLAGNGDYVKGVVGLAKGLRKAESKYPLVVAVLPDVPEDHRKQLVDQGCVVKEIEPVYPPE
24_Cru      -----MAPELTQTTAAKSTVTITKPSPPTIYGGGRAYVTFLAGNGDYVKGVVGLAKGLRKVKSSYPLVVAILPDVPEEHRRILVEQGCIVREIEPVYPPD
25_Ccl      ----MAP-DITPTTITKTTS---LSKTP--SLPKRAYVTFLAGDGDYWKGVVGLVKGLRKAKSKYPLLVAMLPDVPEEHRKILIEQGCFLREIEPVYPPE
26_Ccl      -----MAPPELVQTAVKPAGLGAKPASL----PGRAYVTFLAGNGDYVKGVVGLAKGLRKVKTAYPLVVAVLPDVPEEHRNILESQGCIVREIEPVYPPD
27_Ccl      -----MAP-GVAEDAFSGNGKISSTG-----YSKRAFVTFLAGSGDYVKGVVGLAKGLRNSKSAYPLVVAILPDVPEEHREVLRSQGCIVREIEPIYPPE
28_Egr      -----------MALNVAAAT--KLVKP--ENQPSRAYVTFLAGNGDYVKGVVGLAKGLRKAKSKYPLVVAVLPDVPEEHRKILVDQGCVVREIEPVYPPE
29_Egr      -----------MALNVAAAT--KLVKP--ENQPSRAYVTFLAGNGDYVKGVVGLAKGLRKAKSKYPLVVAVLPDVPEEHRKILVDQGCVVREIEPVYPPE
30_Egr      -----MAP-ELVPASVKPAAK-PVA-------LSRAYVTFLAGTGDYVKGVVGLAKGLRRARSAYPLVVAVLPDVPAEHRLILEEQGCVVREIEPVYPPE
31_Egr      -----MAP-ELVPISVKPADL-PVQAGL----LGRAYLTFLAGNGDYVKGVVGLAKGLRKAKAAYPLVVAVLPDVPEEHRQILESQGCVVREIEPVYPPE
32_Egr      -----------MALNVAAAT--KLMKP--ENQPSRAYVTFLAGNGDYVKGVVGLAKGLRKAKSKYPLVVAVLPDVPEEHRKILVDQGCVVREIEPVYPPE
33_Egr      -----------MAVADAADN--GVVKPKSENQPSCAYVTFLAGNGDYVKGVVGLAKGLRKAESKYPLVVAILPDVPVDHRKILVDQGCIVREIEPVYPPE
34_Atr      -----MAP------EMANGKKNGGYS-------KRAYVTFLAGDGDYIKGAVGLAKGLRKVKTAYPLIVAVLPDVPEEHRRILRSQGCIVREIEPIYPPE
35_Cpa      -----MAP-ELVQNTVKPAAF-VKTARL----PSRAYVTFLAGNGDYVKGVVGLAKGLRKVKSSYPLVVAVLPDVPEEHRRILESQGCIVREIQPVYPPE
36_Gma      -----MAP-ELVPTVVKSSAAFTKPATL----PRRAYVTFLAGNGDYVKGVVGLAKGLRKVKTAYPLVVAVLPDVPEEHRKILESQGCIVREIEPVYPPE
37_Gma      -----MAP-ELVPTV-----AFTKPATL----PRRAYVTFLAGNGDYVKGVVGLAKGLRKVKTAYPLVVAVLPDVPEDHRKILESQGCIVREIEPVYPPE
```

```
38_Gra   -----MAP-ELVQAGVKP-TVLAKPVTL----PKRAYVTFLAGDGDYVKGVVGLAKGLRQVKSAYPLVVAVLPDVPEEHRRILENQGCIVREIEPVYPPE
39_Gra   -----MAPPELVQPSDKT-NVFAKQVTL----RNRAYVTFLAGNGDYVKGVVGLAKGLRKVKSAYPLVVAVLPDVPEEHRRILENQGCIVREIEPVYPPE
40_Zma   -----MSP-ELTGKMAAKAAAAAAA----VKPATRAYVTFLAGDGDYWKGVVGLAKGLRKVGSAYPLVVALLPDVPESHRRILVSQGCILREIEPVYPPE
41_Zma   -----MAPELMTAKMTAKAAAAAAAV----KPATRAYVTFLAGDGDYWKGVVGLAKGLRKVRSAYPLVVAVLPDVPESHRRILVSQGCVVREIEPVYPPE
42_Mac   ---------MAPQMVAGPAAGKGIQAGR---TPSRAYVTFLAGDGDYVKGAVGLSKGLRKVGSAYPLVVAVLPDVPQSHRRLLASQGCIIREIEPVYPPE
43_Mac   --------MASGAPAESLGG--QVKK--------RAFVTFLAGDGDYVKGVVGLAKGLRKVKSAYPLVVAVLPDVPEAHRRLLQAQGCTVREIEPVYPPE
44_Mac   ---------MAPNMKAK---GKGKQAGG---KPSRAYVTFLAGDGDYVKGAVGLAKGLRKVGSAYPLVVAVLPDVPESHRRLLAAQGCIVREIEPVYPPE
45_Kfe   --------MDLSLINTLTAA--SKTASLAKPDQARAYVTFLAGNGDYVKGVVGLAKGLRKVKSKYPLVVAILPDVPAEHRQILVDQGCVVREIVPVYPPE
46_Kfe   -----MAP-ELGSLSVKP----SAIAKP----PSRAYVTFLAGNGDYVKGVVGLVKGLRKVKSAYPLVVAVLPDVPLEHRRILEDQGCIVREIEPVYPPE
47_Kla   --------MDFSLINTLTAA--SKTASLAKPDQARAYVTFLAGNGDYVKGVVGLAKGLRKVKSKYPLVVAILPDVPAEHRQILVDQGCVVREIVPVCPPE
48_Kla   -----MAP-ELGSLSVKP----STIAKS----PSRAYVTFLAGNGDYVKGVVGLVKGLRKVKSAYPLVVAVLPDVPLEHRRILEDQGCIVREIEPVYPPE
49_Kla   -----MAP-ELGSLSVKT----STIAKP----PSRAYVTFLAGNGDYVKGVVGLVKGLRKVKSAYPLIVAVLPDVPLEHRRILEDQGCIVREIEPVYPPE
50_Kla   --------MDFSLINTLTAA--SKTASLAKPDQARAYVTFLAGNGDYVKGVVGLAKGLRKVKSKYPLVVAILPDVPAEHRQILVDQGCVVREIVPVCPPE
51_Osa   -----MAPPQLAGKMTAKAAAAV-------KPATRAYVTFLAGDGDYWKGVVGLAKGLRKVGSAYPLVVAVLPDVPESHRRILISQGCIVREIEPVYPPE
52_Osa   ---------MMGPNVSSEKKALAAA-------KRRAYVTFLAGDGDYWKGVVGLAKGLRRVRSAYPLVVAVLPDVPGEHRRKLVEQGCVVREIQPVYPPE
53_Lus   MAPEIPPTANNAALATKTAAAAGLLKQP--SISKCAYVTFLAGNGDYVKGVVGLAKGLRKVKSKYPLVVAVLPDVPEEHRRILVSQGCIVREIDPVYPPE
54_Mes   -----MAP-ELLQSAMKPVGF-TKPATL----PSRAYVTFLAGNGDYIKGVVGLAKGLRKVKTAYPLVVAVLPDVPEEHRRILESQGCLVREIEPVYPPE
55_Mes   ----MAPNLTTS--TKTTTS---LVKQA--SLSSRAYVTFLAGNGDYVKGVVGLAKGLRKVDSKYPLVVAILPDVPEEHRKILVSQGCIVKEIEPVYPPE
56_Mes   -----MAP-EVPVDVFSGSGKVCAIN-A--GYSKRAYVTFLAGNGDYVKGVVGLAKGLRKVKSAYPLVVAILSDVPDEHRQILKSQGCIVREIEPIYPPE
57_Mdo   -----MAP-ELVPTSVPK-SGFTTPATM----PSRAYVTFLAGNGDYVKGVVGXAKGLRKVKTAYPLVVAVLPDVPEDHRRILESQGCIVREIEPVYPPE
58_Mdo   -----MAP-ELVPTSVPK-SGFTTPATM----PSRAYVTFLAGNGDYVKGVVGXAKGLRKVKTAYPLVVAVLPDVPEDHRRILESQGCIVREIEPVYPPE
59_Mtr   -----MAP-ELVPTAAKSVTGFTKPVTI----PKRAYVTFLAGNGDYVKGVIGLAKGLRKVMTAYPLVVAVLPDVPEEHREMLEAQGCIVREIEPVNPPE
60_Mgu   -----MAPQVKKTDVISGAGKITAVD--------RAYVTFLAGNGDYVKGVVGLAKGLRKVKSAYPLVVAILPDVPEDHRELLRSQGCIVKEIEPIYPPA
61_Mgu   -----MAPVVP-AEVFSSAGKIPANDS------KKGYVTFLAGSGDYWKGVVGLAKGLRKVKSAYPLVVAILPDVPEDHREILRSQGCIVKEIVPIYPPD
62_Fve   -----MAP-ELVPAVPKS-TALTRPATL----PRRAYVTFLAGNGDYVKGVVGLAKGLRKVNTAYPLVVAVLPDVPEDHRRILESQGCIVREIEPVYPPE
63_Csi   -----MAPPELVQTAVKPAGLGAKPASL----PGRAYVTFLAGNGDYVKGVVGLAKGLRKVKTAYPLVVAVLPDVPEEHRNILESQGCIVREIEPVYPPD
64_Oth   -----MAPPQLVGKMTAKKAAA--------KPATRAYVTFLAGDGDYWKGVVGLAKGLRKVGSAYPLVVAVLPDVPESHRRILTSQGCIVREIEPVYPPE
65_Oth   -------------MSTTLSSNHKAAPPV--PMPKRAYVTFLAGDGDYWKGVVGLAKGLRRARSAYPLVVAVLPDVPEEHRRRLAVQGCVVREIKPVYPPE
66_Pha   ----------MGPNMSAYSSKQRAA------PKR-AYVTFLAGDGDYWKGVVGLAKGLRRVRSAYPLVVAVLPDVPEEHRRKLREQGCVVREIQPVYPPE
67_Pha   -----MAP-ELAGKMTAK---AAAAL----KPATRAYVTFLAGDGDYWKGVVGLAKGLRKVGSAYPLVVAVLPDVPESHRRILVSQGCIVREIEPVYPPE
68_Pvi   -------------MSSAYSSKQQAAAK-------RAYVTFLAGDGDYWKGVVGLAKGLRRVRSAYPLVVAVLPDVPEEHRRKLRDQGCVVREIQPVYPPD
69_Pvi   ------------MMSSKQQQQQQQQAAP--NNKRRAYVTFLAGDGDYWKGVVGLAKGLRRVRSAYPLVVAVLPDVPEEHRRKLRGQGCVVREIQPVYPPE
70_Pvi   -----MAP-EIAGKMTAKAAAAAAAV----KPATRAYVTFLAGDGDYWKGVVGLAKGLRKVGSAYPLVVAVLPDVPEAHRRILVSQGCIVREIEPVYPPE
71_Pvi   -----MAP-ELAGKMTAKAA-AAAAV----KPATRAYVTFLAGDGDYWKGVVGLAKGLRKVGSAYPLVVAVLPDVPESHRRILVSQGCIVREIEPVYPPE
72_Stu   ---------MAPNVFGLATKATGLAKAK--SLPSRAYVTFLAGNGDYWKGVVGLVKGLRKAKSAYPLVVACLPDVPEEHRRILINQGCIVREIEPVYPPE
73_Stu   ----------------MAKPVTNGPGPA---TLDRAYVTFLAGNGDYVKGVIGLAKGLRKVKSEYPLVVAVLPDVPAEHRRMLEEQGCVVREIEPVYPPE
74_Ptr   -----MAP-ELVRSALKPAGF-TKLANL----PSRAYVTFLAGDGDYVKGVVGLAKGLRKVKTAYPLIVAVLPDVPEEHRQILESQGCIVREIEPVYPPE
75_Ptr   ----MAP-DITATLANNTNS---LVKQA--SISSCAYVTFLAGDGDYWKGVVGLAKGLRKAKCNYPLVVAILPDVPEEHRKILASQGCIVREIEPVNPPE
76_Ptr   ----MAP-DITTPLANNATT---LVKQA--SISSCAYVTFLAGDGDYWKGVVGLAKGLRKAESKYPLVVAILPDVPEEHRKILVSQGCIVREIEPVHPPE
```

```
77_Ptr   ----MAP-HITTALANSTNS---LVKQA--SLSSCAYVTFLAGDGDYWKGVVGLAKGLRKAKSKYPLVVAILPDVPEEHRMILVSQGCIVREIEPVHPPE
78_Ptr   -----MAP-ELVQAALKPAGF-TKPASL----PSRAYVTFLAGNGDYVKGVVGLAKGLRKVKTAYPLIVAVLPDVPEEHRRILESQGCIVREIEPVYPPE
79_Ppe   -----MAPPEVPVDVFQATSKVSTLSSA--AYSNRAFVTFLAGNGDYVKGVVGLAKGLRKVKSAYPLVVAILPDVPEEHREILRSQGCIVREIEPIYPPE
80_Ppe   -----MAP-ELVPTGVAK-SGFTIPATV----PSRAYVTFLAGNGDYVKGVVGLAKGLRKVKTAYPLVVAVLPDVPQEHRRILESQGCIVREIEPVCPPE
81_Spu   ----MAP-DITTTLANNANT---LVKQA--SISSRAYVTFLAGDGDYWKGVVGLAKGLRKADSKYPLVVAMLPDVPEEHRKILVSQGCIVREIEPVHPPE
82_Spu   ----MAPRIIITTLANTTSS---VVKQA--SLSSCAYVTFLAGDGDYWKGVVGLAKGLRKAKSKYPLVVAMLPDVPEEHRKILVSQGCIVREIEPVHPPE
83_Spu   -----MAP-ELAQAALKPAGF-TKPAGL----PSRAYVTFLAGNGDYVKGVVGLAKGLRKVKSAYPLIVAVLPDVPEEHRRILKSQGCIVREIEPVYPPE
84_Sit   ----------MGPNMSAFSSKQQAS------AKRRAYVTFLAGDGDYWKGVVGLAKGLRKARSAYPLVVAVLPDVPEEHRRKLREQGCVVREIQPVYPPE
85_Sit   -----MAP-ELAGKMTAKAA-AAPAV----KPTTRAYVTFLAGDGDYWKGVVGLAKGLRKVGSAYPLVVAVLPDVPESHRRILVSQGCIVREIEPVYPPE
86_Svi   ----------MGPNMSAFSSKQQAS------AKRRAYVTFLAGDGDYWKGVVGLAKGLRKVRSAYPLVVAVLPDVPEEHRRKLREQGCVVREIQPVYPPE
87_Svi   -----MAP-ELAGKMTAKAA-AAPAV----KPTTRAYVTFLAGDGDYWKGVVGLAKGLRKVGSAYPLVVAVLPDVPESHRRILVSQGCIVREIEPVYPPE
88_Sbi   -----MAP-ELAGKMTAKAAAAAAAAAV-TKPATRAYVTFLAGDGDYWKGVVGLAKGLRKARSAYPLVVAVLPDVPESHRRILVSQGCIVREIEPVYPPE
89_Sbi   ----------MGPNSSALGKQQQAAAA---MAPKRAYVTFLAGDGDYWKGVVGLAKGLRRVGAAYPLVVAVLPDVPEEHRRKLRDQGCVVREIEPVYPPD
90_Sly   -----MAPAIARVTEKMAKPATNGPGPA---TLDRAYVTFLAGNGDYVKGVIGLAKGLRKVKSVYPLVVAVLPDVPPEHRRMLEEQGCIVREIEPVYPPE
91_Sly   ---------MAPNVFGLATKATGLAKAK--SLSSRAYVTFLAGNGDYWKGVVGLVKGLRKAKSAYPLVVACLPDVPEEHRRILINQGCIVREIEPVYPPH
92_Tca   -----MAP-ELVQAAAKP-TAFAKPVTL----PKRAYVTFLAGNGDYVKGVVGLAKGLRKVKSAYPLVVAVLPDVPEEHRRVLENQGCIVREIEPVYPPE
93_Esa   -----MAPELTQTNTAKSAVTITKPPPQ-IYGGDRAYVTFLAGNGDYVKGVVGLAKGLRKVKSAHPLVVAVLPDVPEEHRRLLVEQGCIVREIEPVYPPE
94_Esa   ---------MALEINNKLTE--PVFAATANGGEKRAYVAFLAGNGDFVKGVVALAKGLRKAKSKYPLVVAVLPDVPEDHQKQLVEQGCVIKEIEPVYPPE
95_Tpr   -----MAP-ELIPTATKSVTGFTKSVTI----PKRAYVTFLAGNGDYVKGVIGLAKGLRKVKTAYPLVVAVLPDVPEEHREMLESQGCIVREIEPVYPPE
```

```
                    110       120       130       140       150       160       170       180       190       200
           ....|....|....|....|....|....|....|....|....|....|....|....|....|....|....|....|....|....|....|....|
1_Rco      NQTQFAMAYYVINYSKLRIWEFVEYSKMIYLDGDIQVFDNIDHLFDLPDG-HFYAVMDCFCEKTWSHTPQYKIGYCQQCPDR-VKWPA----K-LGQPPS
2_Rco      NQTQFAMAYYVINYSKLRIWEFVEYSKMIYLDGDIQVFENIDHLFDLQNG-YFYAVMDCFCEKTWSHSPQYKIGYCQQCPDR-VKWPA---EM--GPKPP
3_Rco      NQTQFAMAYYVINYSKLRIWEFVEYSKMIYLDGDIQVFENIDHLFDLQDG-YFYAVMDCFCEKTWSHSPQYKIGYCQQCPDR-VKWPA---EM--GPKPP
4_Aco      NQTQFAMAYYVINYSKLRIWEFVEYKKMIYLDADIQVYDNVDHLFDRPDG-FFYAVMDCFCEKTWSHTPQYKIGYCQQCPDK-VKWPE---HE-LG----
5_Aco      NQTQFAMAYYVINYSKLRIWKFVEYERMIYLDADIQVYENIDHLFDLPHG-YFYAVKDCFCEKTWSHTPQYKIGYCQQCPER-VSWPA---DK-LGPPPP
6_Aco      NHIRFAMAYYVINYSKLRIWNFEEYSKMIYLDADIQVYDNVDHLFDRPDG-YLYAVMDCFCEKTWSHTPQYKIGYCQQCPDK-VAWPA---EM--GPTPP
7_Aly      NQTEFAMAYYVINYSKLRIWKFVEYSKMIYLDGDIQVFDNIDHLFDLPNG-QFYAVMDCFCEKTWSHSPQYKIGYCQQCPDK-VTWPE---AELIGPKPP
8_Aly      NQVEFAMAYYVLNYSKLRIWNFEEYLKMIYLDADIQVFENIDELFDLPDG-YFHAVMDCFCEKTWSHSLQYSIGYCQQCPDK-VTWPE---DM--ESPPP
9_Aly      NQTQFAMAYYVINYSKLRIWKFVEYSKMIYLDGDIQVYENIDHLFDLPDG-YFYAVMDCFCEKTWSHTPQYKIGYCQQCRDK-VQWPK---AE-LGEPPA
10_Aqco    NQTQFAMAYYVINYSKLRIWEFVEYSKMIYLDGDIQVFENIDHLFDLPNG-YFYAVMDCFCEKTWSHTPQYKLGYCQQCPNK-VQWPA---EM--GQPPS
11_Aha     NQVEFAMAYYVLNYSKLRIWNFEEYYKMIYLDADIQVFENIDELFDLPDG-YFHAVMDCFCEKTWSHSLQYSIGYCQQCPER-VTWPE---DM--ESPPP
12_Ath     NQTEFAMAYYVINYSKLRIWEFVEYNKMIYLDGDIQVFDNIDHLFDLPNG-QFYAVMDCFCEKTWSHSPQYKIGYCQQCPDK-VTWPE---AK-LGPKPP
13_Ath     NQTQFAMAYYVINYSKLRIWKFVEYSKMIYLDGDIQVYENIDHLFDLPDG-YLYAVMDCFCEKTWSHTPQYKIRYCQQCPDK-VQWPK---AE-LGEPPA
14_Bst     NETQFAMAYYVINYSKLRIWKFVEYSKMIYLDGDIQVYENIDHLFDLPDG-YFYAVMDCFCEKTWSHTPQYKIGYCQQCPEK-VQWPK---AE-LGEPPA
15_Bst     NQTEFAMAYYVINYSKLRIWKFVEYSKMIYLDGDIQVFDNIDHLFDLPDG-QFYAVMDCFCEKTWSHSPQYKIGYCQQCPDK-VTWPE---AE-LGPKPP
16_Bdi     SQTQFAMAYYVINYSKLRIWEFVEYERMVYLDADIQVFSNIDHLFDLEKG-SFYAVKDCFCEKTWSHTPQFKLGYCQQRPDKNVSWPAD------TPAPP
17_Bdi     NQTQFAMAYYVINYSKLRIWEFVEYERMVYLDADIQVFDNVDELFDLPKG-HFYAVMDCFCEKTWSHTPQYQIGYCQQCPDR-VTWP----AAEMGPPPA
18_Bra     NQTQFAMAYYVINYSKLRIWKFVEYSKMLYLDGDIQVYENIDHLFDLPDG-YFYAVMDCFCEKTWSHTPQYKIGYCQQCPEK-VQWPK---EE-LGEPPS
19_Bra     NQVEFAMAYYVLNYSKLRIWNFEEYSKMMYLDADIQVFDNIDNLFDLPDG-YFHAVMDCFCEKTWSHSPQYSIGYCQQCPEK-VTWPE---DM--ESPPP
20_Bsta    NQTQFAMAYYVINYSKLRIWEFVEYERMVYLDADIQVFGNVDELFDLPKG-HFYAVMDCFCEKTWSHTPQYQIGYCQQCPGR-VTWP----AAEMGPPPA
21_Cgr     NQTEFAMAYYVINYSKLRIWKFVEYSKMIYLDGDIQVFENIDHLFDLPNG-QFYAVMDCFCEKTWSHSPQYKIGYCQQCPDK-VTWPE---AE-LGPKPP
22_Cgr     NQTQFAMAYYVINYSKLRIWKFVEYSKMIYLDGDIQVYENIDHLFDLPDG-YFYAVMDCFCEKTWSHTPQYKIGYCQQCPEK-VQWPK---AE-LGEPPA
23_Cru     NQTEFAMAYYVINYSKLRIWKFVEYSKMIYLDGDIQVFENIDHLFDLPDG-QFYAVMDCFCEKTWSHSPQYKIGYCQQCPDK-VTWPE---AE-LGPKPP
24_Cru     NQTQFAMAYYVINYSKLRIWKFVEYSKMIYLDGDIQVYENIDHLFDLPDG-YFYAVMDCFCEKTWSHTPQYKIGYCQQCPEK-VQWPK---AE-LGEPPA
25_Ccl     NQTQFAMAYYVINYSKLRIWEFVEYSKMIYLDGDIQVFDNIDHLFDLPDG-YFYAVMDCFCEKTWSHTPQFKIGYCQQCPDK-VKWPA---EL--GPKPA
26_Ccl     NQTQYAMAYYVINYSKLRIWEFVEYSKMIYLDGDIQVFENIDHLFDLPDG-YFYAVMDCFCEKTWSKTPQYKIGYCQQCPDR-VRWPA----E-MGEPPA
27_Ccl     NQIQFAMAYYVINYSKLRIWNFEEYSKMIYLDADIQVFENIDHLFDLPDG-FFYAVMDCFCEKTWSHSPQYSIGYCQQRPNK-VTWPA---EM--GSPPP
28_Egr     NQTQFAMAYYVINYSKLRIWEFLEYSKLIYLDGDIQVFDNIDHLFDLPNG-FFYAVMDCFCEKTWSHTPQYQIGYCQQCPDR-IQWPD---HA--GPKPS
29_Egr     NQTQFAMAYYVINYSKLRIWEFLEYSKMIYLDGDIQVFDNIDHLFDLPNG-SFYAVMDCFCEKTWSHTPQFQIGYCQQCPDK-IQWPD---HA--GPKPS
30_Egr     NQTQFAMAYYVINYSKLRIWEFVEYSKMIYLDGDIQVFDNIDHLFDLEDG-YFYAVMDCFCEKTWSHTPQYAIGYCQQCPDR-VRWPD----E-MGPPPA
31_Egr     NQTQFAMAYYVINYSKLRIWEFVEYSKMIYLDGDIQVFENIDHLFDQPDG-YFYAVMDCFCEKTWSHSPQYKVGYCQQSPDR-VKWPA----E-MGEPPA
32_Egr     NQTQFAMAYYVINYSKLRIWEFLEYSKLIYLDGDIQVFDNIDHLFDLPNG-FFYAVMDCFCEKTWSHTPQFQIGYCQQCPDK-IQWPD---HA--GPKPS
33_Egr     NQTQFAMAYYVINYSKLRIWEFVEYDKMIYLDGDIQVFDNIDHLFDLPNS-FFYAVMDCFCEKTWSHTPQYEIGYCQQCPDK-VQWPD---HV--GPKPS
34_Atr     NQTQFAMAYYVINYSKLRIWEYEEYSKMMYLDADIQVFENIDHLFDLPNG-YFYAVMDCFCEKTWSHTPQYKIEYCQQCPDR-VQWPS---HL--GDPPS
35_Cpa     NETQFAMAYYVINYSKLRIWEFVEYSKMIYLDGDIQVFDNIDHLFDLPDV-YFYAVMDCFCEKTWSHSPQYKIGYCQQCPER-VRWPE----E-MGQPPA
36_Gma     NQTQFAMAYYVINYSKLRIWEFVEYSKMIYLDGDIEVYENIDHLFDLPDG-NFYAVMDCFCEKTWSHTPQYKVGYCQQCPEK-VRWPT----E-LGQPPS
37_Gma     NQTQFAMAYYVINYSKLRIWEFVEYTKMIYLDGDIQVYENIDHLFDLPGG-YFYAVMDCFCEKTWSHTPQYKVGYCQQCPEK-VQWPT----E-LGQPPS
```

```
38_Gra   NQTQFAMAYYVINYSKLRIWKFVEYSKMIYLDGDIQVYENIDHLFDLPDG-QFYAVMDCFCEKTWSHTPQFKIGYCQQCPDK-VKWPA----E-MGQPPA
39_Gra   NQTQFAMAYYVINYSKLRIWEFVEYSKMIYLDGDIQVYDNIDHLFDLPDG-HFHAVMDCFCEKTWSHTPQYRIGYCQQCPDK-VKWPA----E-MGNPPS
40_Zma   NQTQFAMAYYVINYSKLRIWEFVEYERMVYLDADIQVFENIDELFELEKG-YFYAVMDCFCEKTWSHTPQYRIGYCQQCPDK-VAWP----TTELGPPPP
41_Zma   NQTQFAMAYYVINYSKLRIWEFVEYERMVYLDADIQVFENIDGLFELEKG-YFYAVMDCFCEKTWSHTPQYRIGYCQQCPDK-VAWPAA--TAELGPPPS
42_Mac   SQTQFAMAYYVINYSKLRIWEFVEYQKMVYLDADIQVYENIDHLFDLPDG-HFYAVMDCFCEKTWSHTPQYKIGYCQQCPGR-VAWPA---DE-LGPPPA
43_Mac   NQVQFAMAYYVINYSKLRIWNFVEYSKIIYLDADIQVYDNIDHLFDMPDG-YFYAVMDCFCEKTWSHSRQYSIGYCQQCPDK-VAWPA---EM--GSPPP
44_Mac   SQTRFAMAYYVINYSKLRLWEFVEYKKMVYLDADIQVFENIDHLFDLPDG-HFYAVMDCFCEKTWSHSPQYKIGYCQQCPDR-VAWPS---DE-LGPPPS
45_Kfe   NQTQFAMAYYVINYSKLRIWEFVEYDKMIYLDGDIQVFENIDHLFDMADG-HFYAVMDCFCERTWSHTPQYSIGYCQQCPDK-VSWPA---SL--GPKPP
46_Kfe   NQTQFAMAYYVINYSKLRIWEFVEYDKMIYLDGDIQVYENIDHLFDLPDG-NFYAVMDCFCEKTWSHTPQYKIGYCQQCPEK-VQWPA----E-LGQPPA
47_Kla   NQTQFAMAYYVTNYSKLRIWEFVEYDKMIYLDGDIQVFENIDHLFDMADG-HFYAVMDCFCERTWSHTPQYSIGYCQQCPDK-VSWPA---SL--GPKPP
48_Kla   NQTQFAMAYYVINYSKLRIWEFVEYDKMIYLDGDIQVYENIDHLFDLPDG-NFYAVMDCFCEKTWSHTPQYKIGYCQQCPEK-VQWPA----E-LGQPPA
49_Kla   NQTQFAMAYYVINYSKLQIWEFVEYDKMIYLDGDIQVYENIDHLFDLPDG-NFYAVMDCFCEMTWSHTPQYKIGYCQQCPEK-VQWPA----E-LGQPPA
50_Kla   NQTQFAMAYYVINYSKLRIWEFVEYDKMIYLDGDIQVFENIDHLFDMADG-HFYAVMDCFCERTWSHTPQYSIGYCQQCPDK-VSWPA---SL--GPKPP
51_Osa   NQTQFAMAYYVINYSKLRIWEFVEYERMVYLDADIQVFDNIDELFELPKG-HFYAVMDCFCEKTWSHTPQYQIGYCQQCPDK-VAWP----TAELGPPPA
52_Osa   SQTQFAMAYYVINYSKLRIWEFVEYERMVYLDADIQVFDNIDHLFDLDKG-AFYAVKDCFCEKTWSHTPQYDIGYCQQRPDE-VAWP----ERELG-PPP
53_Lus   NQTQFAMAYYVINYSKLRIWKFVEYSKMIYLDGDIQVFSNIDHLFDLPDG-RFYAVMDCFCEKTWSHTPQYKIGYCQQRPDK-VKWTP---EL--GPKPA
54_Mes   NQTQFAMAYYVINYSKLRIWEFVEYSKMIYLDGDIQVFENIDHLFDLPDG-YLYAVMDCFCEKTWSHTAQYKIGYCQQCPDR-VKWPA----E-MGQPPS
55_Mes   NQTQFAMAYYVINYSKLRIWEFVEYSKMIYLDGDIQVFENIDHLFELEDG-YFYGVMDCFCEQTWSNSPQYKIGYCQQCPDR-VQWPA---EL--GPKPP
56_Mes   HQIQFAMAYYVINYSKLRIWNFEEYSKMMYLDADIQVFENIDHLFDAPDG-YFYAVMDCFCEKTWSHSAQYSIGYCQQCPDR-VTWPT---DM--GSPPP
57_Mdo   NQTQFAMAYYVINYSKLRIWEFVEYEKMIYLDGDIEXYDNIDHLFDLPDG-HFYAVMDCFCEKTWSHTPQYQIGYCQQCPEK-VQWPT---SE-LGPPPS
58_Mdo   NQTQFAMAYYVINYSKLRIWEFVEYEKMIYLDGDIEXYDNIDHLFDLPDG-HFYAVMDCFCEKTWSHTPQYQIGYCQQCPEK-VQWPT---SE-LGPPPS
59_Mtr   NQTQFAMAYYVINYSKLRIWEFVEYSKMIYLDGDIQVYENIDHLFDLPDG-HFYAVMDCFCERTWSHTPQYKIGYCQQCPEK-VHWPK----E-MGQPPS
60_Mgu   NQIQFAMAYYVINYSKLRIWNFLEYSKMVYLDADIQVYENIDHLLDAPDG-YFYAVMDCFCEKTWSHSRQFSVGYCQQCPNK-VTWPA---EM--GPPPP
61_Mgu   NQIQFAMAYYVINYSKLRIWDFEEYSKMVYLDADIQVYENIDHLLDTPNG-YFYAVMDCFCEKTWSHSKQYSIGYCQQCPNK-VTWPA---GM--GAPPP
62_Fve   NQTQFAMAYYVINYSKLRIWEFVEYDKMIYLDGDIQVYDNIDHLFDLPDG-NFYAVMDCFCEKTWSHTPQYKIGYCQQCPER-VKWDF----E-LGPPPS
63_Csi   NQTQYAMAYYVINYSKLRIWEFVEYSKMIYLDGDIQVFENIDHLFDLPDG-YFYAVMDCFCEKTWSKTPQYKIGYCQQCPDR-VRWPA----E-MGEPPA
64_Oth   NQTQFAMAYYVINYSKLRIWEFVEYERMVYLDADIQVYDNIDELFELEKG-HFYAVMDCFCEKTWSHTPQYKIGYCQQCPDR-VAWP----TAELGPPPA
65_Oth   NQTQFAMAYYVINYSKLRIWEFVEYERMVYLDADIQVFDNIDELFDLEKGGHFYAVMDCFCEKTWSHTPQYKVGYCQQCPDR-VTWP----ERELGCPPP
66_Pha   SQTQFAMAYYVINYSKLRIWEFVEYERMVYLDADIQVYENIDHLLDLEKG-RFYAVMDCFCEKTWSHTPQYKIGYCQQCPER-VSWPEQ--EQELG-PPP
67_Pha   NQTQFAMAYYVINYSKLRIWEFVEYERMVYLDADIQVFENIDELFELEKG-HFYAVMDCFCEKTWSHTPQYKIGYCQQCPDK-VAWP----AAELGPPPS
68_Pvi   SQTQFAMAYYVINYSKLRIWEFVEYERMVYLDADIQVFDNIDELFELDKG-RFYAVMDCFCEKTWSHTPQYKIGYCQQCPEP-VAWP----AAELG-PPP
69_Pvi   SQTQFAMAYYVINYSKLRIWELVEYERMVYLDADIQVLDNIDHLLDLDKG-RFYAVMDCFCEKTWSHTPQYKIGYCQQCPER-VPWP----EQQLG-PPP
70_Pvi   NQTQFAMAYYVINYSKLRIWEFVEYERMVYLDADIQVFDNIDELFELEKG-HFYAVMDCFCEKTWSHTPQYKIGYCQQCPDK-VAWP----TAELGPPPA
71_Pvi   NQTQFAMAYYVINYSKLRIWEFVEYERMVYLDADIQVFENIDELFELEKG-HFYAVMDCFCEKTWSHTPQYKIGYCQQCPDK-VAWP----TAELGPPPP
72_Stu   NQTQFAMAYYVINYSKLRIWEFVEYSKMIYLDGDIQVFDNIDHLFDLPDG-YFYAVMDCFCEKTWSHTPQYKVGYCQQCPDK-VQWTQ---DL--GPKPS
73_Stu   NQTQFAMAYYVINYSKLRMWEFVEYKKMIYLDGDIQVYDNIDHLFDLADG-YFYAVMDCFCEKTWSHTPQYKIGYCQQCPDR-IKWPS---DE-LGQPPS
74_Ptr   NQTQFAMAYYVINYSKLRIWEFVEYSKMIYLDGDIQVYDNIDHLFDLPDG-RFYAVMDCFCEKTWSHTPQYKIGYCQQCPDK-VNWPA----E-MGQPPS
75_Ptr   NQTQFAMAYYVINYSKLRIWEFVEYSKMIYLDGDIQVFDSIDHLFDMPDG-YFYAAMDCFCEKTWSNSPQYKIGYCQQCPDK-VHWPA---EM--GPKPP
76_Ptr   NQTRFAMPYYVINYSKLRIWEFVEYSKMIYLDGDIQVFDNIDHLFDMPDG-CFYAVMDCFCEKTWSNSPQYKIGYCQQCPDK-VQWPA---EM--GPKPP
```

```
77_Ptr    NQTRFAMPYYVINYSKLRIWEFVEYSKMIYLDGDIQVFDNIDHLFDMPDG-YFYAVMDCFCEKTWSNSPQYKIGYCQQCPDK-VQWPA---EM--GPKPP
78_Ptr    NQTQFAMAYYVINYSKLRIWEFVEYSKMIYLDGDIQVYDNIDHLFDLPDG-HFYAVMDCFCEKTWSHTPQYKIGYCQQCPDK-VNWPA----E-MGQPPS
79_Ppe    NQIKFAMAYYVINYSKLRIWNFEEYSKMIYLDADIQVYENIDHLFDTPDG-YFYAVMDCFCEKTWSHSPQYKVGYCQQCPDK-VSWPA---EL--GSPPP
80_Ppe    NQTQFAMAYYVINYSKLRIWEFVEYSKMIYLDGDIQVYDNIDHLFDLPDG-HFYAVMDCFCEKTWSHTPQYKIGYCQQCPEK-VQWPA---SE-LGPPPS
81_Spu    NQTQFALAYYVINYSKLRIWEFVEYSKMIYLDGDIQVFDNIDHLFDMPGG-FIHAVMDCFCEKTWSISPQYKIGYCQQCPDK-VRWPA---EM--GPRPP
82_Spu    NQTRFAMPYYVINYSKLRIWEFVEYSKMIYLDGDIQVFDNIDHLFDMPGG-HFYAVMDCFCEKTWSNSPQYKIGYCQQCPDK-VRWPA---EM--GPKPP
83_Spu    NQTQFAMAYYVINYSKLRIWEFVEYTKMIYLDGDIQVYDNIDHLFDLTDG-HFYAVMDCFCEKTWSHTPQYKIGYCQQCPGK-VNWPD----E-MGQPPS
84_Sit    SQTQFAMAYYVINYSKLRIWEFVEYERMVYLDADIQVYENIDHLFDLEKG-RFYAVMDCFCEKTWSHTPQYKIGYCQQCPDK-VTWP----EHELG-PPP
85_Sit    NQTQFAMAYYVINYSKLRIWEFVEYERMVYLDADIQVFENIDELFELEKG-SFYAVMDCFCEKTWSHTPQYKIGYCQQCPDR-VAWP----TAELGPPPA
86_Svi    NQTQFAMAYYVINYSKLRIWEFVEYERMVYLDADIQVYENIDHLFDLEKG-RFYAVMDCFCEKTWSHTPQYKIGYCQQCPDK-VTWP----EHELG-PPP
87_Svi    NQTQFAMAYYVINYSKLRIWEFVEYERMVYLDADIQVFENIDELFELEKG-SFYAVMDCFCEKTWSHTPQYKIGYCQQCPDK-VAWP----TAELGPPPA
88_Sbi    NQTQFAMAYYVINYSKLRIWEFVEYERMVYLDADIQVFENVDELFELEKG-YFYAVMDCFCEKTWSHTPQYKIGYCQQCPDK-VAWPA---TAELGPPPA
89_Sbi    SQTQFAMAYYVINYSKLRIWELVEYERMVYLDADIQVYSNIDHLFDLEKG-KFHAVMDCFCEKTWSHTPQYKIGYCQQCPER-VAWPEQEQEQELG-PPP
90_Sly    NQTQFAMAYYVINYSKLRIWEFVEYKKLIYLDGDIQVYDNIDHLFDLPDG-YLYAVMDCFCEKTWSHTPQYKIGYCQQCPDK-VKWPS---ED-LGQPPS
91_Sly    NQTQFAMAYYVINYSKLRIWEFVEYSKMIYLDGDIQVFDNIDHLFDLPDG-YFYAVMDCFCEKTWSHTPQYKVGYCQQCPDK-VQWTE---DL--GPKPS
92_Tca    NQTQFAMAYYVINYSKLRIWEFVEYSRMIYLDGDIQVFDNIDHLFDLPDG-HCYAVMDCFCEKTWSHTQQYKIGYCQQCPDK-VKWPA----E-MGQPPS
93_Esa    NQTQFAMAYYVINYSKLRIWKFVEYSKLIYLDGDIQVYDNIDHLFDLDDG-YFYAVMDCFCEKTWSHTPQYKIGYCQQCPEK-VQWPK---AE-LGEPPA
94_Esa    NQTDFAMAYYVINYSKLRIWKFVEYSKMIYLDGDIQVFENIDHLFDLPNG-HFYAAKDCFCEKTWSHTPQYKIGYCQQCPDK-VTWPE---AE-LGPKPP
95_Tpr    NQTQFAMAYYVINYSKLRIWEFLEYSKMIYLDGDIQVYDNIDHLFDLPDG-YFYGVMDCFCEKTWSHTPQYKIGYCQQCPQK-VQWPK----Q-MGQPPS
```

```
                  210       220       230       240       250       260       270       280       290       300
         ....|....|....|....|....|....|....|....|....|....|....|....|....|....|....|....|....|....|....|....|
1_Rco    -LYFNAGMFVFEPSISTYHDLLKTVQITPPTPFAEQDFLNMYFRDIYKPIPVVYNLVLAMLWRHPENVE---LDKVKVVHYCAAGSKPWRYTGKEENMER
2_Rco    -LYFNAGMFVFEPSLSTYDDLLNTVKLTPPTPFAEQDFLNMFFKDIYRPIPPIYNLVLALLWRHPENIE---FEKVKVVHYCAAGSKPWRYTGKEDNMDR
3_Rco    -LYFNAGMFVFEPSLPTYDDLLNTVKLTPPTPFAEQDFLNMFFKDIYRPIPPIYNLVLALLWRHPENIE---LEKVKVVHYCAAGSKPWRYTGKEENMDR
4_Aco    -------MFVSEPSLATAKSLLETLEITPPTPFAEQDFLNMFFKDIYKPIPLVYNLVLAMLWRHPDNVE---LDKVKVVHYCAAGSKPWRYTGKEANMDR
5_Aco    -LYFNAGMFVSEPDTAIADSLLAALKVAPTTPFAEQDFLNMFFKDIYKPIPLVYNLVLAMLWRHPENVE---LDKVKVVHYCAA---PWRFTGKEPNMDR
6_Aco    -LYFNAGMFVFEPGREICDRLLETLRVVPVTPFAEQDFLNMFFEKIYKPIPLVYNLVLAMLWRHPENVE---LDNVKVVHYCAAGSKPWRYTGKEANMDR
7_Aly    -LYFNAGMFVYEPNLSTYHSLLETVKVVPPTLFAEQDFLNMYFKDIYKPIPPVYNLVLAMLWRHPENIE---LDQVKVVHYCAAGAKPWRFTGEEENMDR
8_Aly    PLYFNAGMFVFEPSPLTYESLLQTLEITPPSPFAEQDFLNMFFEKVYKPIPLVYNLVLAMLWRHPENVE---LEKVKVVHYCAAGSKPWRYTGEEANMDR
9_Aly    -LYFNAGMFLFEPNLETYEDLLRTLKITPPTPFAEQDFLNMYFKKIYKPIPLVYNLVLAMLWRHPENVE---LGKVKVVHYCAAGSKPWRYTGKEANMDR
10_Aqco  -LYFNAGMFVYEPSLSTYDDLLKQVKTTPPTPFAEQDFLNMFFKDIYKPIPLVYNLVLAMLWRHPENVE---LEKVKVVHYCAAGSKPWRYTGKEENMDR
11_Aha   PLYFNAGMFVFEPSPLTYESLLHTLEITPPSPFAEQDFLNMFFEKVYKPIPLVYNLVLAMLWRHPENVE---LEKVKVVHYCAAGSKPWRYTGEEANMDR
12_Ath   -LYFNAGMFVYEPNLSTYHNLLETVKIVPPTLFAEQDFLNMYFKDIYKPIPPVYNLVLAMLWRHPENIE---LDQVKVVHYCAAGAKPWRFTGEEENMDR
13_Ath   -LYFNAGMFLYEPNLETYEDLLRTLKITPPTPFAEQDFLNMYFKKIYKPIPLVYNLVLAMLWRHPENVE---LGKVKVVHYCAAGSKPWRYTGKEANMER
14_Bst   -LYFNAGMFLFEPNLETYEDLLRTLKITPPTPFAEQDFLNMYFKKIYKPIPLVYNLVLAMLWRHPENVE---LDKVKVVHYCAAGSKPWRYTGKEANMDR
15_Bst   -LYFNAGMFVYEPNLSTYHNLLETLKVVPPTLFAEQDFLNMYFKDIYKPIPSAYNLVLAMLWRHPENIE---LDQVKVVHYCAAGAKPWRFTGEEENMDR
16_Bdi   PLYFNAGMFVHEPSMATARALLEKLVVTDPTPFAEQDFLNVFFRDAYKPIPLVYNLVLAMLWRHPENVE---LDAVKVVHYCAAGSKPWRFTGEEENMER
17_Bdi   -LYFNAGMFVHEPSMATAKALLDTLRVSPTTPFAEQDFLNVFFREQYKPIPLVYNLVLAMLWRHPENVQ---LAKVKAVHYCAAGSKPWRFTGKEANMDR
18_Bra   -LYFNAGMFVFEPGLDTYEDLLRTLKITPPTPFAEQDFLNMYFEKIYKPIPLVYNLVLAMLWRHPENVE---LDKVKVVHYCAAGSKPWRYTGKEANMER
19_Bra   PLYFNAGMFLFEPSPLTYESLLQTLETTPPSPFAEQDFLNMFFEKVYKPIPLVYNLVLAMLWRHPENVD---LEKVKVVHYCAAGSKPWRYTGEEANMDR
20_Bsta  -LYFNAGMFVHEPSMATAKALLDSLRVSPTTPFAEQDFLNVFFREQYKPIPLMYNLVLAMLWRHPENVQ---LEKVKAVHYCAAGSKPWRFTGKEANMDR
21_Cgr   -LYFNAGMFVYEPNLSTYHNLLETLQVVPPTLFAEQDFLNMYFKDIYKPIPSAYNLVLAMLWRHPENIE---LDQVKVVHYCAAGAKPWRFTGEEENMDR
22_Cgr   -LYFNAGMFLFEPNLETYEDLLKTLQITPPTPFAEQDFLNMYFKKIYKPIPLVYNLVLAMLWRHPENVE---LDKVKVVHYCAAGSKPWRYTGKEANMDR
23_Cru   -LYFNAGMFVYEPNLSTYHNLLETLQVVPPTLFAEQDFLNMYFKDIYKPIPSAYNLVLAMLWRHPENIE---LDQVKVVHYCAAGAKPWRFTGEEENMDR
24_Cru   -LYFNAGMFLFEPNLETYEDLLKTLQITPPTPFAEQDFLNMYFKKIYKPIPLVYNLVLAMLWRHPENVE---LDKVKVVHYCAAGSKPWRYTGKEANMDR
25_Ccl   -LYFNAGMFVFEPSLSTYHDLLETVQITTPTSFAEQDFLNMYFRDIYRPIPPIYNLVVAMLWRHPENVE---ADKAKVVHYCAAGSKPWRFTGKEENMQR
26_Ccl   -LYFNAGMFVFEPSISTYHDLLETVKVTPPTTFAEQDFLNMYFKHIYKPIPLVYNLVLAMLWRHPENVE---LDKVKVVHYCAAGSKPWRFTGEEENMQR
27_Ccl   -LYFNAGMFVFEPSRLTYENLLQTLKITTPTPFAEQDFLNMFFQKTYKPIPLAYNLVVAMLWRHPENVE---LEKVKAVHYCAAGSKPWRYTGKEVNMDR
28_Egr   -LYFNAGMFVYEPNLKTYRDLLEKLKITPPTSFAEQDFLNEYFKDIYKPIPNVYNFVLAMLWRHPKNVE---LNKVKVVHYCAAGSKPWRYTGEEQNMDR
29_Egr   -LYFNAGMFVYEPNLKTYRDLLEKLKITPPTSFAEQDFLNEYFKDIYKPIPNVYNFVLAMLWRHPENVE---LDKVKVVHYCAAGSKPWRYTGEEQNMDR
30_Egr   -LYFNAGMFVYEPSLATYHDLLSAVKVTPPTSFAEQDFLNMYFRNVYKPIPLAYNLILAMLWRHPENVE---LEKVKVVHYCAAGSKPWRYTGKEENMQR
31_Egr   -LYFNAGMFVFEPSIATYVDLLDTLKITAPTSFAEQDYLNMYFRNIYKPIPLDYNLVLAMLWRHPENVD---LEKVKVVHYCAAGSKPWRYTGKEENMQR
32_Egr   -LYFNAGMFVYEPNLKTYRDLLEKLKITPPTSFAEQDFLNEYFKDIYKPIPNVYNFVLAMLWRHPENVE---LDKVKVVHYCAAGSKPWRYTGEEQNMDR
33_Egr   -LYFNAGMFVYEPNLYTYHDMLETLKITPPTPFAEQDFLNMYFKDIYKPIPNVYNLVLAMLWRHPENVE---LDKVKVVHYCAAGSKPWRYTGEEQHMDR
34_Atr   -LYFNAGMFVFEPSKLTYENLLQTLQVTTPTPFAEQDFLNMFFKDVYKPIPLVYNLVMAMLWRHPENVD---LETVKVVHYCAAGSKPWRYTGKEENMQR
35_Cpa   -LYFNAGMFVFEPSLYTYQDLLSTVKVTPTTPFAEQDFLNMYFKDIYKPIPLVYNLVLAMLWRHPEKVE---LDKVKVVHYCATGSKPWRYTGKEENMQR
36_Gma   -LYFNAGMFVFEPNIATYHDLLKTVQVTTPTSFAEQDFLNMYFKDIYKPIPLNYNLVLAMLWRHPENVK---LDQVKVVHYCAAGSKPWRYTGKEENMQR
37_Gma   -LYFNAGMFVFEPSIATYHDLLKTVQVTTPTSFAEQDFLNMYFKDIYKPIPLNYNLVLAMLWRHPENVK---LDQVKVVHYCAAGSKPWRYTGKEENMQR
```

```
38_Gra   -LYFNAGMFVFEPSLTTYENLLATLKSTPPTPFAEQDFLNMYFKDIYKPIPLVYNLVLAMLWRHPDNVE---LDKVKVVHYCAAGSKPWRYTGKEENMQR
39_Gra   -LYFNAGMFVFEPSLVTYESLLKTLKITQPTPFAEQDFLNMFFKDIYKPIPLIYNLVLAMLWRHPENVE---LEKVKVVHYCAAGSKPWRYTGKEANMQR
40_Zma   -LYFNAGMFAHEPSMATAKALLDTLRVTPPTPFAEQDFLNMFFRDQYRPIPNVYNLVLAMLWRHPENVQ---LEKVKVVHYCAAGSKPWRFTGKEANMDR
41_Zma   -LYFNAGMFVHEPSVATAKALLDTLRVTPPTPFAEQDFLNMFFRDQYRPIPNVYNLVLAMLWRHPENVQ---LEKVKVVHYCAAGSKPWRFTGKEANMDR
42_Mac   -LYFNAGMFVHEPSLATCESLLSTLKVTPPTPFAEQDFLNMFFKDIYKPIPLIYNLVLAMLWRHPENVE---LEKVKVVHYCAAGSKPWRYTGKEANMDR
43_Mac   -LYFNAGMFVSEPARPTYDGLLETLMTTPPTPFAEQDFLNMYFEKIYKPIPLIYNLVLAMLWRHPENVE---LENVKVVHYCAAGSKPWRYTGKEANMDR
44_Mac   -LYFNAGMFVHEPSLATSESLLAALKITCPTPFAEQDFLNMFFKDIYKPIPLVYNLVLAMLWRHPENVE---LEKVKVVHYCAAGSKPWRFTGKEANMDR
45_Kfe   -LYFNAGMFVYEPSLTTYHDLLDTLMVAPTTPFAEQDFLNTYFSKIYKPIPNIYNLVLAMLWRHPENVE---LGKVKVVHYCAAGSKPWRYTGEEANMDR
46_Kfe   -LYFNAGMFVFEPSIATYHDLLKTLMITPPTPFAEQDFLNMYFRNIYKPIPLVYNLVLAMLWRHPENVE---LEKVKVVHYCAAGSKPWRYTGKEPNMDR
47_Kla   -LYFNAGMFVYDPSLTTYHDLLDTLMVAPTTPFAEQDFLNTYFSKIYKPIPNTYNLVLAMLWRHPENVE---LGKVKVVHYCAAGSKPWRYTGEEANMDR
48_Kla   -LYFNAGMFVFEPSIATYHDLLKTLMITPPTPFAEQDFLNMYFRNIYKPIPLVYNLVLAMLWRHPENVD---LEKVKVVHYCAAGSKPWRYTGKEPNMDR
49_Kla   -LYFNAGMFVFEPSIATYHDLLKTLMITPPTPFAEQDFLNMYFRNIYKPIPLVYNLVLAMLWRHPENVE---LEKVKVVHYCAAGSKPWRYTGKEPNMDR
50_Kla   -LYFNAGMFVYEPSLTTYHDLLDTLMVAPTTPFAEQDFLNTYFSKIYKPIPNTYNLVLAMLWRHPENVE---LGKVKVVHYCAAGSKPWRYTGEEANMDR
51_Osa   -LYFNAGMFVHEPSMATAKSLLDTLRVTTPTPFAEQDFLNMFFREQYKPIPLIYNLVLAMLWRHPENVQ---LEKVKVVHYCAAGSKPWRYTGKEANMDR
52_Osa   PLYFNAGMFVHEPGLGTAKDLLDALVVTPPTPFAEQDFLNMFFREQYKPIPNVYNLVLAMLWRHPENVD---LDQVKVVHYCAAGSKPWRFTGKEENMNR
53_Lus   -LYFNAGMFVYEPSLAIYDKLLETLKVAPTTPFAEQDFLNSFFKDIYRPIPSYYNLVLAMLWRHPENVE---LEKVKVVHYCAAGSKPWRYTGEEENMDR
54_Mes   -LYFNAGMFVFEPSVSTYHDLLKTVKITPPTSFAEQDFLNMYFKDIYKPIPLVYNLVLAMLWRHPENVE---LDEVKVVHYCAAGSKPWRYTGKEENMQR
55_Mes   -LYFNAGMFVFEPNLSTYHDLLTTVKVTTPTLFAEQDFLNMFFRDKYRPIPPIYNLVLAMLWRHPENVE---LDKLKVVHYCAAGSKPWRYTGKDANMDR
56_Mes   -LYFNAGMFVFEPSRLTYDNLLHTLKITPPTPFAEQDFLNMFFEKTYKPLPLVYNLVLAMLWRHPENVE---VEKVKVAHYCAAGSKPWRYTGKEANMDR
57_Mdo   -LYFNAGMFVFEPGLETYHDLLRTLRVTPPTPFAEQDFLNMYFRKIYKPIPLVYNLVLAMLWRHPENVE---LDKVKVVHYCAAGSKPWRYTGKEENMQR
58_Mdo   -LYFNAGMFVFEPGLETYHDLLRTLRVTPPTPFAEQDFLNMYFRKIYKPIPLVYNLVLAMLWRHPENVE---LDKVKVVHYCAAGSKPWRYTGKEENMQR
59_Mtr   -LYFNAGMFLFEPSIDTYHDLLKTLKVTPPTPFAEQDFLNMYFKDIYKPIPFVYNLVLAMLWRHPENVE---LHKVKVVHYCAAGSKPWRYTGKEENMQR
60_Mgu   -LYFNAGMFVYEPNKTTYEALLETLQITPPTPFAEQDFLNMFFEKTYQPIPLVYNLVLAMLWRHPENVE---LDKVKVVHYCAAGSKPWRYTGEEANMDR
61_Mgu   -LYFNAGMFVYEPNKTTYENLLETLKITTPTPFAEQDFLNMFFEKTYKPIPLIYNLVLAMLWRHPENVE---LEKVQVVHYCAAGSKPWRYTGEEANMDR
62_Fve   -LYFNAGMFVFEPSVLTYQDLLKTLRVAPTTPFAEQDFLNMYFRDIYKPIPNVYNLVLAMLWRHPENVQ---LEKVKVVHYCAAGSKPWRYTGKEENMER
63_Csi   -LYFNAGMFVFEPSISTYHDLLETVKVTPPTTFAEQDFLNMYFKHIYKPIPLVYNLVLAMLWRHPENVE---LDKVKVVHYCAAGSKPWRFTGEEENMQR
64_Oth   -LYFNAGMFVHEPSMETAKALLDTLRVTPPTPFAEQDFLNMFFRDQYKPIPNIYNLVLAMLWRHPENVQ---LEKIKVVHYCAAGSKPWRYTGKEANMDR
65_Oth   PLYFNAGMFVHEPSTETARALLDTLRVTPPTPFAEQDFLNVFFRDVYRPIPNVYNLVLAMLWRHPENVE---LDKVKVVHYCAAGSKPWRYTGEEPNMDR
66_Pha   PPYFNAGMFVHEPSLGTAKDLLDALVVTPPTPFAEQDFLNMFFRDVYSPIPPVYNLVLAMLWRHPDKVE---LGKVKVVHYCAAGSKPWRYTGQEPNMER
67_Pha   -LYFNAGMFVHEPSMATAKALLDTLRVTPPTPFAEQDFLNMFFREQYKPIPLVYNLVLAMLWRHPENVQ---LEKVKVVHYCAAGSKPWRYTGKEANMDR
68_Pvi   PLYFNAGMFVHEPSLGTARDLLDALVATPPTPFAEQDFLNVFFRDVYTPIPPAYNLVLAMLWRHPDKMPE--LGKVKVVHYCAAGSKPWRYTGQEPNMER
69_Pvi   PPYFNAGMFVHEPSLGTAKELLDALVATPPTPFAEQDFLNMFFRDVYTPIPPVYNLVLAMLWRHPDKVEP--LAEVKVVHYCAAGSKPWRFTGQEPNMER
70_Pvi   -LYFNAGMFVHEPSMATAKALLETLRVTPPTPFAEQDFLNMFFREQYKPIPLVYNLVLAMLWRHPENVQ---LEKVKVVHYCAAGSKPWRYTGKEANMDR
71_Pvi   -LYFNAGMFVHEPSMATAKALLDTLRVTPPTPFAEQDFLNMFFREQYKPIPLVYNLVLAMLWRHPENVQ---LEKVKVVHYCAAGSKPWRYTGKEANMDR
72_Stu   -LYFNAGMFVYEPNLSTYDDLLKTVKVTPPTPFAEQDFLNMYFRDVYKPIPNNYNLVLAMLWRHPENVD---LEKVKVVHYCAAGSKPWRYTGKEENMDR
73_Stu   -LYFNAGMFVFEPSLRTYQDLLKKLQITPPTPFAEQDFLNMYFKNIYRPIPLVYNLVLAMLWRHPENVE---LDKVKVVHYCAAGSKPWRYTGKEENMER
74_Ptr   -PYFNAGMCVFEPSIATYHDLLKTLKVTPPTPFAEQDFLNMYFKDIYTPIPLVYNLVLAMLWRHPENVE---LDRVKVVHYCAAGSKPWRYTGKEENMQR
75_Ptr   -LYFNAGMFVYEPNLSTYHDLLETLKVTTPTLFAEQDFLNMFFRDVYKPIPSDYNLVLALLWRHPENIN---LDKVKVVHYCAAGSKPWRYTGKEDNMDR
76_Ptr   -LYFNAGMFVYEPNLSTYHDLLETLKITSPTLFAEQDFLNMFFRDVYKPIPSDYNLVLAMLWRHPENIN---LDKVKVVHYCAAGSKPWRYTGKEENMDR
```

| | |
|---|---|
| 77_Ptr | -LYFNAGMFVYEPNLSTYHDLLETVKVTSPTLFAEQDFLNMFFRDVYKPIPSDYNLVLAMLWRHPENIN---LDKVKVVHYCAAGSKPWRFTGKEENMDR |
| 78_Ptr | -LYFNAGMFVFEPSISTYHDLLKTLKVTPPTPFAEQDFLNMYFKDIYKPIPLVYNLVLAMLWRHPDNVE---LDKVKVVHYCAAGSKPWRYTGKEENMQR |
| 79_Ppe | -LYFNAGMFVFEPSRLTYDSLLQTLQIVPPTPFAEQDFLNMFFQKTYKPIPLVYNLVLAMLWRHPENVE---LDKVNVVHYCAAGSKPWRYTGKEANMDR |
| 80_Ppe | -LYFNAGMFVFEPDLETYHDLLNTLKVTPPTPFAEQDYLNMFFRKIYKPIPLAYNLVLAMLWRHPENVE---LDKVKVVHYCAAGSKPWRYTGKEENMER |
| 81_Spu | -LYFNAGMFVYEPKLSTYHDLLETLKVTTPTLFAEQDFLNMFFRDVYKPIPSDYNLVLAMLWRHPENIY---LDKVKAVHYCAAGSKPWRYTGKEENMDR |
| 82_Spu | -LYFNAGMFVYEPNLSTYHDLLGTLKTTSPTLFAEQDFLNMFFRDVYEPIPSDYNLVLAMLWRHPENIN---LDKVKVVHYCAAGSKPWRFTGKEENMDR |
| 83_Spu | -LYFNAGMFVFEPSISTYHDLLKTLKVTPPTPFAEQDFLNMYFKNIYKPIPLVYNLVLAMLWRHPENVE---LDKVKVVHYCAAGSKPWRYTGKEENMQR |
| 84_Sit | PRYFNAGMFVHEPSLGTAKDLLDALVVTPPTPFAEQDFLNMFFRDVYEPIPPVYNLVLAMLWRHPENVKP--LDKVKVVHYCAAGSKPWRYTGEEANMDR |
| 85_Sit | -LYFNAGMFVHEPSMATAKALLDTLRVTPPTPFAEQDFLNMFFREQYKPIPNVYNLVLAMLWRHPENVQ---LEKVKVVHYCAAGSKPWRYTGKEANMDR |
| 86_Svi | PRYFNAGMFVHEPSLGTAKDLLDALVVTPPTPFAEQDFLNMFFRDVYEPIPPVYNLVLAMLWRHPENVKP--LDKVKVVHYCAAGSKPWRYTGEEANMDR |
| 87_Svi | -LYFNAGMFVHEPSMATAKALLDTLRVTPPTPFAEQDFLNMFFREQYKPIPNVYNLVLAMLWRHPENVQ---LEKVKVVHYCAAGSKPWRYTGKEPNMDR |
| 88_Sbi | -LYFNAGMFVHEPSMATAKALLDTLRVTPPTPFAEQDFLNMFFRDQYRPIPNVYNLVLAMLWRHPENVQ---LEKVKAVHYCAAGSKPWRFTGKEPNMDR |
| 89_Sbi | PLYFNAGMFVHEPSLRTAKDLLDALVVTPPTPFAEQDFLNLFFRDVYSPIPPVYNLVLAMLWRHPDKLKVVRLDEVKVVHYCAAGSKPWRYTGKEPNMDR |
| 90_Sly | -LYFNAGMFVFEPSLRTYHDLLKKLQITPPTPFAEQDFLNMYFKNIYRPIPLVYNLVLAMLWRHPENVE---LDKVKVVHYCAAGSKPWRYTGKEENMER |
| 91_Sly | -LYFNAGMFVYEPSLSTYDDLLKTLKVTPPTPFAEQDFLNMYFRDVYKPIPNDYNLVLAMLWRHPENVD---LEKVKVVHYCAAGSKPWRYTGKEENMDR |
| 92_Tca | -LYFNAGMFVFEPSLFTYKNLLETLKITPPTPFAEQDLLNMFFRDIYKPIPLLYNLVLAMLWRHPENVE---LDKVKVVHYCAAGSKPWRWTGKEENMHR |
| 93_Esa | -LYFNAGMFVFEPDLNTYEDLLRTLKITPPTPFAEQDFLNMYFKKIYKPIPLVYNLVLAMLWRHPENVE---LDKVKVVHYCAAGSKPWRYTGKETNMER |
| 94_Esa | -LYFNAGMFVYEPNLYTYQNLLETLKVVPPTSFAEQDFLNMYFKDIYTPIPGVYNLVMAMLWRHPENVE---LEQVKVVHYCAAGSKPWRFTGKEENMER |
| 95_Tpr | -LYFNAGMFLFEPSIETYHDLLNTLQVTPPTPFAEQDFLNMYFKDIYKPIPLVYNLVLAMLWRHPENVD---IHNVKVVHYCAAGSKPWRYTGKEDNMQR |

```
                  310       320       330       340       350       360       370       380       390
         ....|....|....|....|....|....|....|....|....|....|....|....|....|....|....|....|....|....|..
1_Rco    EDIKMVVKKWWDVYNDESLDYKKQ-------------PAADG--DAEPMN-LQPFIA-ALYEAG---AVQYVTAPSAA--------------
2_Rco    EDIKMLVKKWWDIYEDESLDYKNT-V-------------AATGGATE-GE-LQPFLA-ALSEAG---VVHYVTAPSAA--------------
3_Rco    EDIKTLVKKWWDIYEDESLDYKNTAA-------------AATGGATE-AG-LQPLLA-AMSEAS---EVHYITAPSAA--------------
4_Aco    EDIKMLVKKWWDIYNDESLDYKGPAVA-------AVDAISVAVEAEPEPNALQKPFLAALSEAG---KVQYITAPSAA--------------
5_Aco    EDIKMLVSKWWDIYNDDSLDYKG----------------DVTVATEAEANGVKPPLKAALSKAG---VVHYIKAPSAAIIESTDGYLGSLLS
6_Aco    EDIKTLVAKWLDIYNDDSLDYVAAEN-------------AVLQET----ACSLSPIVTVVTETS---NNSYIPTPSAA--------------
7_Aly    EDINMLVKKWWDIYNDESLDYKNVVI-------------GDGHKKQ---QTLQQFIE-ALSEAG---VLQYVKAPSAA--------------
8_Aly    EDIKMLVDKWWDVYNDESLDFKSIIP-------------ADVEET-----VTKSSILASVLEPE---IT-YFPAPSAA--------------
9_Aly    EDIKMLVKKWWDIYNDESLDYKKPVA----------VVGT----EADPVN-LKPFIT-ALTEAG---RVNYVTAPSAA--------------
10_Aqco  EDIKMLVKKWWDIYDDASLDYKRS-V-------------APSIPETG--ENLQPFLA-ALSEAG---VVHYVTAPSAA--------------
11_Aha   EDIKMLVDKWWDVYNDESLDFKSKIP-------------ADAEET-----VTKSSILASVLEPE---IT-YFPAPSAA--------------
12_Ath   EDIKMLVKKWWDIYNDESLDYKNVVI-------------GDSHKKQ---QTLQQFIE-ALSEAG---ALQYVKAPSAA--------------
13_Ath   EDIKMLVKKWWDIYDDESLDYKKPVT----------VVDT----EVDLVN-LKPFIT-ALTEAG---RLNYVTAPSAA--------------
14_Bst   EDIKMLVNKWWDIYNDDSLDYMKPVV----------IVDT----EADLTN-LKPFIT-ALTEAG---RVNYVTAPSAA--------------
15_Bst   EDIKLLVKKWWDIYNDESLDYKNIVI-------------ADSHKKQ---QTFQQFIE-ALSEAG---VLQYVKAPSAA--------------
16_Bdi   EDVKMLVKKWWDVYNDDSLDFDDSYKELK-------KKSEAGGVVFDHEAGGKPARRGATAMADGAGAVKYSNTPSAA--------------
17_Bdi   EDIKVLVKKWWDIYNDESLDFKG------------LP----ADAD-ELEAAAKKPIRAALAEAG---TVKYITAPSAA--------------
18_Bra   EDIKMLVNKWWDIYNDDSLDYKKSVG----------DLVE----ESDVVN-LKPFIS-ALTEAG---PVKYVTAPSAA--------------
19_Bra   EDIKMLVDKWWDVYNDESLDFKSKIP-------------VDVEET-----VTKSTILASVLEPE---MT-YIPAPSAA--------------
20_Bsta  EDIKVLVKKWWDIYNDESLDFKG------------LP----ADAD-ELEAAAKKPIRAALAEAG---TVKYITAPSAA--------------
21_Cgr   EDIKMLVKKWWDIYNDESLDYKN-II-------------GDGYKKQ---QTFQQFVE-ALSEAG---VLQYVKAPSAA--------------
22_Cgr   EDIKMLVNKWWDIYKDDSLDYKKPLG----------VAET----EADLVN-LKPFIT-ALTEVG---RVNYVTAPSAA--------------
23_Cru   EDIKMLVKKWWDIYNDESLDYKN-II-------------GDGYKKQ---QTFQQFVE-ALSEAG---VLQYVKAPSAA--------------
24_Cru   EDIKMLVNKWWDIYKDDSLDYKKPLG----------VAET----EADLVN-LKPFIT-ALTDVG---RVNYVTAPSAA--------------
25_Ccl   EDIKMLVQKWWDIFNDESLDYKNYVP-------------PDEDAEDE-AARAQPFLM-ALSEAG---VVYYVAAPSAA--------------
26_Ccl   EDVKMLVKKWWDIYNDESLDYKK--------------PSADG--NAGSVN-LQPFID-ALSDAA---AVQFVTAPSAA--------------
27_Ccl   EDVKMLVTKWWDIYNDESLDFDADQN-------------PSAPEEE---TFSKSKIIAPMPEPA---IS-YIPAPSAA--------------
28_Egr   EDIKMLVKKWWDIYDDESLDYRNIVA-------------RDEAAKR---ANLERFLA-ALSEAG---VVPYVPAPSAA--------------
29_Egr   EDIKMLVKKWWDIYDDESLDYRNIVA-------------RDEAAKR---ANLERFLA-ALSEAG---VVPYVPAPSAA--------------
30_Egr   EDIKMLVKKWWDIYEDESLDYKKKAA-----------VAAAAQGEAEPVN-MEPFFA-ALSEAG---EVQYVTAPSAA--------------
31_Egr   EDIKMLVQKWRDIYNDESLDYKG-TE-----------VEAVEK----AVT-VEPFIA-SLSEVPTA-PVQYVTAPSAA--------------
32_Egr   EDIKMLVKKWWDIYDDESLDYRNIVA-------------RDEAVKR---ANLERFLA-ALFEAG---VVPYVPAPSAA--------------
33_Egr   DDIKMLVKNWWDIYDDESLDYKNIVA-------------RDAAAKQ---TKWDRFLA-ALAEAG---AFRFVTAPSAA--------------
34_Atr   EDIKVLVKKWGDIYNDESLDFKGEPL-------------LAPDLQQPQQPAGHAFYVAAMAEAA---PVQFIPAPSAA--------------
35_Cpa   EDIKMLVKKWWDIYNDESLDYKP--------------VLGGED-----IH-LEPFIA-ALSEAG---QVQYVTAPSAA--------------
36_Gma   EDIKMLVKKWWDIYNDASLDYKPLMN------------ASEAPAADGVD--IEQFVQ-ALSEVG---HVQYVTAPSAA--------------
37_Gma   EDIKMLVKKWWDIYNDASLDYKPLMS------------ASEAPAPDGVDN-IEPFVQ-ALSEVG---HVQYVTAPSAA--------------
```

```
38_Gra   EDVKMLVQKWWDIYNDESLDYEKP--------------TIGEGQAAESVN-MQPFLV-ALSEAG---AVQYVTAPSAA--------------
39_Gra   EDIKMLVQKWWDIYNDESLDYRT---------------SAAEG-GTETVN-LQPFLV-ALSEVG---AVHFVAAPSAA--------------
40_Zma   EDIKTLVNKWWDIYNDEALDFKG-LP---------LSP---ADADDEVEAVAKKPLRAALAEAG---TVKYVTAPSAA--------------
41_Zma   EDINALVNKWWDIYNDETLDLKG------------LPSLS-PDDDDEVEAVAKKPLRAALAEAG---TVKYVTAPSAA--------------
42_Mac   EDIKILVKRWWDVYNDTSLDYKG------------AAATEGVVVPAAAEEERKQPLLAALSEAG---AIKYVTAPSAA--------------
43_Mac   EDIKMLVAKWWGIYNDKSLDF-AAED-------------AVPEGE----TCLPSSIMVAMSENN---IN-YISAPSAA--------------
44_Mac   KDINVLVKRWWDVYHDESLDYKG------------PAVTS------------KQPLRGALPEAG---PVKYVTAPSAA--------------
45_Kfe   EDIKMLVKKWWDVYTDESLDYWNHNV-------------VAAAAALPVNGEIPLLNA-ALKKAG---ADQYVTAPSAA--------------
46_Kfe   EDIKMLVKKWWDIYNDETLDLKRQQQQ-PPQAVASAIPVPADEVSPEPVS-MQPFIE-ALTQAG---AVQFVAAPSAA--------------
47_Kla   EDIKMLVKKWWDVYTDESLDYWNHN--------------AAAAAALPVNGEIPLLNA-ALKKAG---ADQYVTAPSAA--------------
48_Kla   EDIKMLVKKWWDIYNDETLDLKRQQQQ-PPQAVPSAVPVPADEVSPEPVS-MQPFIE-ALTQAG---AVQFVAAPSAA--------------
49_Kla   EDIKMLVKKWWDIYNDETLDLKRQQQQLPPQAAPSAVPVPADEVSPEPVS-MQPFIE-ALTQAG---AVQFVAAPSAA--------------
50_Kla   EDIKMLVKKWWDVYTDESLDYWNHNV-------------VAAAAALPVNGEIPLLNA-ALKKAG---ADQYVTAPSAA--------------
51_Osa   EDIKMLVKKWWDVYNDGSLDFKG------------LPPIAAADDADEVEAAAKKPLRAALAEAR---TVKYVTAPSAA--------------
52_Osa   EDIKMLVKRWWDIYNDESLDYKEEED---------------NAD------EASQPMRTALAEAG---AVKYFPAPSAA--------------
53_Lus   EDIKMLVKKWWDVYDDESLDYWNWVE-------------ATEVTVEA-VK-MQPFLA-ALSEAG---VVHYVTAPSAA--------------
54_Mes   EDIKMLVKKWWDIYNDESLDYKK--------------PAGDG--DAEPVK-LQPFIA-ALSEAG---ALQYVTAPSAA--------------
55_Mes   EDIKMLVKKWWDIYNDESLDYKNTVG-------------AAGGGGIE-GG-LQLPLA-ALSEGG---VVHYISAPSAA--------------
56_Mes   EDIKMLVKKWWEIYNDESLDLKPENS-------------VAVAEDE---TLTRTSIMASMPEPA---IS-YIPAPSAA--------------
57_Mdo   EDIKMLVKKWWDIYNDESLDYKKAAG---GAGAGGVVGPEAASGGERVN--MRPFID-ALSEA----AVQYVTAPSAA--------------
58_Mdo   EDIKMLVKKWWDIYNDESLDYKKAAG---GAWAGGVVGPEAASGGEGVN--MQPFID-ALSEA----AVQYVTAPSAA--------------
59_Mtr   EDIRMLVKKWWDIYNDSSLDYNKNLS------------GSGEVQTNGFE--IEPFVQ-ALSEVG---RVQYVTAPSAA--------------
60_Mgu   EDIKMLVKKWWDVYDDESLDFKPDE---------------AFSKP---------SIMASMPEPA---VS-YIPAPSAA--------------
61_Mgu   EDIKMLVKKWWDVYDDASLDFKAEDS-------------VAGSRP---------SIMASMPEPA---VS-YIPAPSAA--------------
62_Fve   EDIKMLVQKWWEIYNDESLDYKKPQS---------STVVAAVEAEQAVD--MQPFIE-ALSEAG---AFQYVNVPSAA--------------
63_Csi   EDVKMLVKKWWDIYNDESLDYKK--------------PSADG--NAGSVN-LQPFID-ALSDAA---AVQFVTAPSAA--------------
64_Oth   EDIKVLVKKWWDIYDDETLDFKG------------LPDMP----ADEVEAAAKKPLRAALAEAG---TVKYVTAPSAA--------------
65_Oth   DDIKMLVNKWWDIFNDETLDYKGPPLP-------------LHEDDDHAGAAGEPLRRQALSEAG---AAKYFPAPSAA--------------
66_Pha   EDIKMLVKKWWDVFNDESLDYKGPAV---------------DG-------EARQPLRQALAEAG---AAKYFPAPSAA--------------
67_Pha   EDIKMLVKKWWDIYNDETLDFKA------------LP--A-TDAD-EVEAVAKKPIRAALAEAG---TVKYVTAPSAA--------------
68_Pvi   DDIKMLVTKWWDVFNDDTLDYKGD------------------EP----------LL-QALAEAG---AANYFPAPSAA--------------
69_Pvi   QDIKMLVARWWDVFNDDSLDYKGDDA---------------DQP----------LLRQALAEAG---AANYFPAPSAA--------------
70_Pvi   EDIKMLVKKWWDIYNDESLDFKG------------LP-AA-DDDD-EVEAVAKKPIRAALAEAG---TVKYVTAPSAA--------------
71_Pvi   EDIKTLVKKWWDIYNDESLDFKG------------LPAAA-ADAD-EVEAVAKKPIRAALAEAG---TVKYVTAPSAA--------------
72_Stu   EDIKMLIKKWWDIYDDVSLDYKNSNV-------------VMTAVDGE--VEAEKFMA-ALSEAG---VVHYITAPSAA--------------
73_Stu   EDIKLLVKKWWDIYNDESLDYKRSVG----------MNQVNVIGAAGAVNQLQPLIAAAMSQAG---AVKYVTAPSAA--------------
74_Ptr   EDIKMLVKKWWDIYSDESLDSKK--------------LVADCTTDAEPVN-LQPFIA-ALSEAG---AVQYVTAPSAA--------------
75_Ptr   EDIKMLVNKWWDIYHDESLDYKNTVV-------------AAAGA-----E-VQPFLA-ALSEAG---IAHYITAPSAA--------------
76_Ptr   EDIKMLVQKWWDIYNDESLDHKN--T-------------VVASS-GS-E--LQPILE-ALYEAG---VDLHFTAPSAA--------------
```

```
77_Ptr    EDIKMVVNKWWDIYQDESLDYKNTVA-------------AAAAS-AG-AE-LHPFLA-ALSEAG---VVHYVTAPSAA--------------
78_Ptr    EDIKMLVEKWWGIYNDESLDYMK--------------FVADG-IDAEPVN-LQSFIA-ALYEAG---AVQYVTAPSAA--------------
79_Ppe    EDIKMLVAKWWEVYNDETLDFKAEN--------------PADAEEE---AFARSSIMASMPEPA---IS-YIPAPSAA--------------
80_Ppe    EDIKMLVKKWWDIYNDKSLDYKKPSAP--GARAGTRAAVVGGAEGEGVN--MQPFIE-ALSEAG---VVQYVTAPSAA--------------
81_Spu    EDIKMLVQKWWDVYNDELLDYKNSTT-------------VAASS-GS-DQ-LQPILE-ALFEDGG--GHSCFTAPSAA--------------
82_Spu    EDIKMVVNKWWDIYHDETLDYKNTAA-------------AAAAAEAE-AE-LQPFLA-ALSEAG---AVHYVTAPSAA--------------
83_Spu    EDIKTLVEKWWGIYNDESLDYMK--------------SMADC-IDAEPVN-LQPFIA-ALSEAG---AVQYVTAPSAA--------------
84_Sit    EDIKMLVSKWWDIFNDESLDYKGPAV---------------DDDGAEVVDQAREPLRQALAEAG---AAKFFPAPSAA--------------
85_Sit    EDIKMLVNKWWNIYNDESLDFKG------------LPALP-ADAD-EVEAVAKKPIRAALAEAG---TMKYVTAPSAA--------------
86_Svi    EDIKMLVSKWWDIFNDESLDYKGPAV---------------DDDGAEVVDQAREPLRQALAEAG---AAKFFPAPSAA--------------
87_Svi    EDIKMLVNKWWNIYNDETLDFKG------------LPALP-ADAD-EVEAVAKKPIRAALAEAG---TMKYVTAPSAA--------------
88_Sbi    EDIKMLVKKWWDIYNDETLDFKGLLP---------LPPAD-ADADDEVEAVAKKPLRAALAEAG---TVKYVTAPSAA--------------
89_Sbi    DDIKALVAKWWHIFDDQTLDYNGGEA---------------AAD------QASLPLRQALAQAG---AVKYFPAPSAA--------------
90_Sly    EDIKLLVKKWWDIYNDESLDYNRSVG----------MNQVNVIGAAGAVNQLQPLIAAAMSQAS---AVKYVTAPSAA--------------
91_Sly    EDIKMLIKKWWDIYDDESLDYKNSNV-------------VMNAVDGE--VEAQKIME-ALSEAG---VVHYITAPSAA--------------
92_Tca    EDIKMLVQKWWDIYNDESLDYKKP--------------TVAEG-AAEPVN-PQPFLA-ALSEAG---AVQYVTAPSAA--------------
93_Esa    EDIKMLVNKWWDIYNDDSLDYTKSVA----------DLER----DSDMVN-LKPFIT-ALTEAG---RVKYVTAPSAA--------------
94_Esa    EDIKVLVKKWWDIYNDKSLDYKNVIG-------------EKGGNPD---LHKKQLVE-ALSEAG---VLQYVKAPSAA--------------
95_Tpr    EDIKMLVKKWWDVYNDSSLDYNKNLS------------GNSEAHTNGVE--NEPFVQ-ALSEVG---RVQYVTAPSAA--------------
```
