## Supplemental Tables S1-S3 for "Biochemical characterization of recombinant UDP-sugar pyrophosphorylase and galactinol synthase from *Brachypodium distachyon*"

**Supplemental Table S1**

Protein sequences used for phylogenetic tree reconstruction. Sequences were number-coded for clarity in the phylogenetic tree (see Fig. 2). All sequences were obtained from the Phytozome v12.1 server.

| Code | Organism | Family | Transcript name |
|---|---|---|---|
| 1_Rco | *R. communis* | Euphorbiaceae | 30076.m004605 |
| 2_Rco | *R. communis* | Euphorbiaceae | 30128.m008945 |
| 3_Rco | *R. communis* | Euphorbiaceae | 30128.m008946 |
| 4_Aco | *A. comosus* | Bromeliaceae | Aco001744.1 |
| 5_Aco | *A. comosus* | Bromeliaceae | Aco002583.1 |
| 6_Aco | *A. comosus* | Bromeliaceae | Aco005386.1 |
| 7_Aly | *A. lyrata* | Brassicaceae | AL1G66490.t1 |
| 8_Aly | *A. lyrata* | Brassicaceae | AL2G16220.t1 |
| 9_Aly | *A. lyrata* | Brassicaceae | AL4G46730.t1 |
| 10_Aqco | *A. coerulea* | Ranunculaceae | Aqcoe2G257100.1.p |
| 11_Aha | *A. halleri* | Brassicaceae | Araha.32566s0001.1.p |
| 12_Ath | *A. thaliana* | Brassicaceae | AT1G56600.1 |
| 13_Ath | *A. thaliana* | Brassicaceae | AT2G47180.1 |
| 14_Bst | *B. stricta* | Brassicaceae | Bostr.25993s0046.1.p |
| 15_Bst | *B. stricta* | Brassicaceae | Bostr.8169s0004.1.p |
| 16_Bdi | *B. distachyon* | Poaceae | Bradi1g17200.1.p |
| 17_Bdi | *B. distachyon* | Poaceae | Bradi1g64120.1.p |
| 18_Bra | *B. rapa FPsc* | Brassicaceae | Brara.D02832.1.p |
| 19_Bra | *B. rapa FPsc* | Brassicaceae | Brara.I01595.1.p |
| 20_Bst | *B. stacei* | Poaceae | Brast02G160300.1.p |
| 21_Cgr | *C. grandiflora* | Brassicaceae | Cagra.11315s0002.1.p |

| | | | |
|---|---|---|---|
| 22_Cgr | *C. grandiflora* | Brassicaceae | Cagra.6400s0002.1.p |
| 23_Cru | *C. rubella* | Brassicaceae | Carubv10020539m |
| 24_Cru | *C. rubella* | Brassicaceae | Carubv10025703m |
| 25_Ccl | *C. clementina* | Rutaceae | Ciclev10001308m |
| 26_Ccl | *C. clementina* | Rutaceae | Ciclev10021027m |
| 27_Ccl | *C. clementina* | Rutaceae | Ciclev10032043m |
| 28_Egr | *E. grandis* | Myrtaceae | Eucgr.B01791.1.p |
| 29_Egr | *E. grandis* | Myrtaceae | Eucgr.B01793.1.p |
| 30_Egr | *E. grandis* | Myrtaceae | Eucgr.H00906.1.p |
| 31_Egr | *E. grandis* | Myrtaceae | Eucgr.K03563.1.p |
| 32_Egr | *E. grandis* | Myrtaceae | Eucgr.L00234.1.p |
| 33_Egr | *E. grandis* | Myrtaceae | Eucgr.L00245.1.p |
| 34_Atr | *A. trichopoda* | Amborellaceae | evm_27.model.AmTr_v1.0_scaffold00003.201 |
| 35_Cpa | *C. papaya* | Caricaceae | evm.model.supercontig_145.23 |
| 36_Gma | *G. max* | Fabaceae | Glyma.03G222000.1.p |
| 37_Gma | *G. max* | Fabaceae | Glyma.19G219100.1.p |
| 38_Gra | *G. raimondii* | Malvaceae | Gorai.001G145800.1 |
| 39_Gra | *G. raimondii* | Malvaceae | Gorai.007G120900.1 |
| 40_Zma | *Z. mays* | Poaceae | GRMZM2G165919_P01 |
| 41_Zma | *Z. mays* | Poaceae | GRMZM5G872256_P02 |
| 42_Mac | *M. acuminata* | Musaceae | GSMUA_Achr11P16140_001 |
| 43_Mac | *M. acuminata* | Musaceae | GSMUA_Achr6P27280_001 |
| 44_Mac | *M. acuminata* | Musaceae | GSMUA_Achr8P10810_001 |
| 45_Kfe | *K. fedtschenkoi* | Crassulaceae | Kaladp0057s0103.1.p |
| 46_Kfe | *K. fedtschenkoi* | Crassulaceae | Kaladp0085s0102.1.p |
| 47_Kla | *K. laxiflora* | Campanulaceae | Kalax.0267s0048.1.p |

| | | | |
|---|---|---|---|
| 48_Kla | *K. laxiflora* | Campanulaceae | Kalax.0285s0036.1.p |
| 49_Kla | *K. laxiflora* | Campanulaceae | Kalax.0397s0014.1.p |
| 50_Kla | *K. laxiflora* | Campanulaceae | Kalax.0803s0006.1.p |
| 51_Osa | *O. sativa* | Poaceae | LOC_Os03g20120.1 |
| 52_Osa | *O. sativa* | Poaceae | LOC_Os07g48830.1 |
| 53_Lus | *L. usitatissimum* | Linaceae | Lus10031434 |
| 54_Mes | *M. esculenta* | Euphorbiaceae | Manes.05G012000.1.p |
| 55_Mes | *M. esculenta* | Euphorbiaceae | Manes.09G179100.1.p |
| 56_Mes | *M. esculenta* | Euphorbiaceae | Manes.13G029000.1.p |
| 57_Mdo | *M. domestica* | Rosaceae | MDP0000446914 |
| 58_Mdo | *M. domestica* | Rosaceae | MDP0000466683 |
| 59_Mtr | *M. truncatula* | Fabaceae | Medtr7g109920.1 |
| 60_Mgu | *M. guttatus* | Phrymaceae | Migut.E00169.1.p |
| 61_Mgu | *M. guttatus* | Phrymaceae | Migut.L01457.1.p |
| 62_Fve | *F. vesca* | Rosaceae | mrna24544.1-v1.0-hybrid |
| 63_Csi | *C. sinensis* | Rutaceae | orange1.1g019647m |
| 64_Oth | *O. thomaeum* | Poaceae | Oropetium_20150105_01144A |
| 65_Oth | *O. thomaeum* | Poaceae | Oropetium_20150105_10155A |
| 66_Pha | *P. hallii* | Poaceae | Pahal.B05006.1 |
| 67_Pha | *P. hallii* | Poaceae | Pahal.D02454.1 |
| 68_Pvi | *P. virgatum* | Poaceae | Pavir.Ba00293.1.p |
| 69_Pvi | *P. virgatum* | Poaceae | Pavir.Bb03717.1.p |
| 70_Pvi | *P. virgatum* | Poaceae | Pavir.J07018.1.p |
| 71_Pvi | *P. virgatum* | Poaceae | Pavir.J40731.1.p |
| 72_Stu | *S. tuberosum* | Solanaceae | PGSC0003DMP400006276 |
| 73_Stu | *S. tuberosum* | Solanaceae | PGSC0003DMP400009269 |

| | | | |
|---|---|---|---|
| 74_Ptr | *P. trichocarpa* | Salicaceae | Potri.002G191600.1 |
| 75_Ptr | *P. trichocarpa* | Salicaceae | Potri.005G006800.1 |
| 76_Ptr | *P. trichocarpa* | Salicaceae | Potri.013G005800.1 |
| 77_Ptr | *P. trichocarpa* | Salicaceae | Potri.013G005900.1 |
| 78_Ptr | *P. trichocarpa* | Salicaceae | Potri.014G116800.1 |
| 79_Ppe | *P. persica* | Rosaceae | Prupe.1G251600.1.p |
| 80_Ppe | *P. persica* | Rosaceae | Prupe.3G005100.1.p |
| 81_Spu | *S. purpurea* | Salicaceae | SapurV1A.0088s0080.1.p |
| 82_Spu | *S. purpurea* | Salicaceae | SapurV1A.0088s0090.1.p |
| 83_Spu | *S. purpurea* | Salicaceae | SapurV1A.0294s0080.1.p |
| 84_Sit | *S. italica* | Poaceae | Seita.2G438500.1.p |
| 85_Sit | *S. italica* | Poaceae | Seita.9G424800.1.p |
| 86_Svi | *S. viridis* | Poaceae | Sevir.2G450900.1.p |
| 87_Svi | *S. viridis* | Poaceae | Sevir.9G429000.1.p |
| 88_Sbi | *S. bicolor* | Poaceae | Sobic.001G391300.1.p |
| 89_Sbi | *S. bicolor* | Poaceae | Sobic.002G423600.1.p |
| 90_Sly | *S. lycopersicum* | Solanaceae | Solyc01g079170.2.1 |
| 91_Sly | *S. lycopersicum* | Solanaceae | Solyc02g084980.2.1 |
| 92_Tca | *T. cacao* | Malvaceae | Thecc1EG004881t1 |
| 93_Esa | *E. salsugineum* | Brassicaceae | Thhalv10001530m |
| 94_Esa | *E. salsugineum* | Brassicaceae | Thhalv10024122m |
| 95_Tpr | *T. pratense* | Fabaceae | Tp57577_TGAC_v2_mRNA4274 |

**Supplemental Table S2**

Comparative analysis of kinetic parameters from different plant USPPases. ND, not determined.

| **Enzyme** | **Organism** | **Source** | **MM (kDa)** | $S_{0.5}$ **Gal1P (mM)** | $S_{0.5}$ $UTP_{Gal1P}$ **(mM)** | $V_{max}$ **(U mg$^{-1}$)** | $S_{0.5}$ **Glc1P (mM)** | $S_{0.5}$ $UTP_{Glc1P}$ **(mM)** | $V_{max}$ **(U mg$^{-1}$)** | $S_{0.5}$ **GlcA1P (mM)** | $S_{0.5}$ $UTP_{GlcA1P}$ **(mM)** | $V_{max}$ **(U mg$^{-1}$)** | **Reference** |
|---|---|---|---|---|---|---|---|---|---|---|---|---|---|
| UGGPase | *Cucumis melo* | Recombinant | 68 | 0.43 | ND | 714 | 0.27 | ND | 222 | ND | ND | ND | Dai *et al* ., 2006 |
| PsUSP | *Pisum sativum* | Recombinant | 67 | 0.58 | ND | 161 | 0.34 | 0.048 | 106 | 0.48 | ND | 66 | Kotake *et al* ., 2004 |
| AtUSP | *Arabidospis thaliana* | Recombinant | 70 | 0.27 | ND | 86.9 | 0.23 | ND | 84.3 | 0.094 | ND | 51.4 | Kotake *et al* ., 2007 |
| AtUSP | *Arabidospis thaliana* | Recombinant | 69 | ND | ND | 24.6 | 0.42 | 0.19 | 41.7 | 0.13 | 0.14 | 32.1 | Litterer *et al* ., 2006 |
| USP1 | *Glycine max* | Recombinant | 71 | ND | ND | 112.91 | 0.23 | 0.19 | 110.48 | 0.14 | 0.15 | 97 | Litterer *et al* ., 2006 |
| PdUSPase | *Populus deltoides* | Recombinant | 51.5 | ND | ND | ND | 1.49 | 1.98 | 23.4 | ND | ND | ND | Kim *et al* ., 2013 |
| BdiUSPPase | *Brachypodium distachyon* | Recombinant | 68.17 | 0.25 | 0.109 | 1024 | 0.28 | 0.11 | 757 | 1.4 | 0.14 | 183.1 | This work |

**Supplemental Table S3**

Comparative analysis of kinetic parameters from different plant GolSases. ND, not determined; a, nM $min^{-1}$; b, U $gFW^{-1}$; c, mM $min^{-1}$ $mg^{-1}$.

| Enzyme | Organism | Source | MM (kDa) | $S_{0.5}$ UDP-Gal (mM) | $S_{0.5}$ *myo*-inositol (mM) | $V_{max}$ (U $mg^{-1}$) | Reference |
|---|---|---|---|---|---|---|---|
| BvGolS1 | *Beta vulgaris* | Recombinant | 39.1 | 0.79 | 4.76 | NR | Kito *et al*., 2018 |
| BvGolS1 | *Beta vulgaris* | Root | 39.1 | ND | ND | $4.70 \times 10^{-6}$ | Kito *et al*., 2018 |
| BvGolS1 | *Beta vulgaris* | Leaf | 39.1 | ND | ND | $0.38 \times 10^{-6}$ | Kito *et al*., 2018 |
| CsGolS1 | *Camellia sinensis* | Recombinant | 38.97 | ND | ND | 0.19 | Zhou *et al*., 2017 |
| CsGolS2 | *Camellia sinensis* | Recombinant | 38.69 | ND | ND | 0.08 | Zhou *et al*., 2017 |
| CsGolS3 | *Camellia sinensis* | Recombinant | 37.8 | ND | ND | 0.3 | Zhou *et al*., 2017 |
| PaXgGolS1 | *Hybrid poplar* | Recombinant | ND | 0.8 | ND | 657.5[a] | Unda *et al*., 2011 |
| PaXgGolS2 | *Hybrid poplar* | Recombinant | ND | 0.65 | ND | 1245[a] | Unda *et al*., 2011 |
| GS | *Phaseolous vulgaris* | Kidney bean | 38 | 0.4 | 4.5 | 8.75 | Liu *et al*., 1995 |
| GS | *Curcubita pepo* | Leaf | 36 | ND | ND | 32.1 | Liu *et al*., 1995 |
| GS | *Curcubita pepo* | Leaf | 42 | 1.8 | 6.5 | 23.3 | Smith *et al*., 1991 |
| GolSase | *Cucumis sativus* | Leaf | ND | 0.16 | 4 | 0.81 | Handley *et al*., 1982 |
| GS | *Ajuga reptans* | Leaf | ND | 0.53 | 6.4 | 0.303[b] | Bachmann *et al*., 1994 |
| BnGolS | *Brassica napus* | Recombinant | 39.4 | 0.401 | 4.96 | 73[c] | Li *et al*., 2011 |
| BdiGolSase1 | *Brachypodium distachyon* | Recombinant | 37.9 | 0.06 | 2.5 | 28.6 | This work |
